# Serotonergic reinforcement of a complete swallowing circuit

**DOI:** 10.1101/2023.05.26.542464

**Authors:** Andreas Schoofs, Anton Miroschnikow, Philipp Schlegel, Ingo Zinke, Casey M Schneider-Mizell, Albert Cardona, Michael J Pankratz

## Abstract

How the body interacts with the brain to perform vital life functions such as feeding is one of the fundamental questions in physiology and neuroscience. Here, we use a whole-animal scanning transmission electron microscopy dataset of *Drosophila* to map out the neuronal circuits that connect the entire enteric nervous system to the brain via the insect vagus nerve at synaptic resolution. This revealed a periphery-brain feedback loop in which Piezo-expressing mechanosensory neurons sense food intake and convey that information onto serotonergic neurons within the brain. These serotonergic neurons integrate the interoceptive information with external and central inputs, and in turn stabilize rhythmic activity of serotonin receptor 7 expressing peripheral motor neurons that drive swallowing. Strikingly, the very same motor neurons also share an efference copy of their activity with the aforementioned mechanosensory neurons, thereby closing the motor-sensory-modulatory loop. Our analysis elucidates an elemental, albeit surprisingly complex reinforcement circuit in which rhythmic motor patterns are stabilized through afferent signaling to central serotonergic neurons upon completion of a rewarding action. The circuit motif is constructed to allow the distinction between self-generated action and those in response to the environment.

## INTRODUCTION

One of the most vital functions of any organism is to take in nutrients from the environment. Feeding in animals entails extensive interaction with multitudinous sensory signals offered by nature, as well as with complex physiological signals provided by the internal organs. This behavior can be seen as having different modules that form a chain of events, each of which requires distinct sets of actions to optimally exploit the range of possible choices based on external offering and internal state over time (Swanson, 2012; Tinbergen, 1951). These include, for example, the central pattern generators (CPGs) that underlie rhythmic motor patterns for pharyngeal pumping, chewing and swallowing (Delcomyn, 1980; Harris-Warrick et al., 1992). Distinct CPGs underlie these steps, but they must be interconnected to move the food from one station to another in a coordinated manner. Moreover, a particular motor pattern should be reinforced when it successfully fulfills a biological need. How this is accomplished at the neuronal circuit level is not well understood.

The various sensory and metabolic signals that underlie feeding behavior, and the neuronal circuits underlying these processes, have been a topic of much interest in different organisms (Friedman, 2019; Miroschnikow et al., 2020; Münch et al., 2020; Sternson and Eiselt, 2017). These include the role of enteric neurons in nutrient sensing and the neuronal pathways by which interoceptive signals are transmitted to the brain (Hadjieconomou et al., 2020; Kim et al., 2020, 2021; Min et al., 2021; Wang et al., 2020). However there is an issue that has attracted much discussion on a conceptual and physiological level, but for which little is known on a neuronal circuit level, namely, how does an organism distinguish its own actions due to its intrinsic spontaneous or rhythmic CPGs versus actions that actually fulfill a biological need. For example, the CPGs for the swallowing reflex in insects and mammals are located in the subesophageal zone (SEZ) and the brainstem, respectively (Jean, 2001; Schoofs et al., 2014a), and swallowing movements can occur in the absence of food. What mechanisms exist by which the body registers that real food has been swallowed and communicates this information to the nervous system such that the action should be strengthened and continued? This issue has been addressed within different behavioral and physiological contexts, and concepts have been proposed on how this might be manifested in neuronal terms (Jékely et al., 2021; Straka et al., 2018; Von Holst, 1953). For example, the terms “exafference” and “reafference” have been used to distinguish afferent signals that come from the environment (exafference, e.g., swallowing of food) from those due to self-generated actions (reafference, e.g., swallowing movements without food). Efference copy and corollary discharge have also been used to refer to the process where the motor system sends signals to the sensory system concerning its own activity. However, the precise nature of the neuronal architecture has not been fully elucidated at circuit and cellular level due to lack of experimental tools. A necessary step to addressing this issue is to completely characterize the architecture of a functional circuit that connects the brain and the peripheral organs at single cell resolution.

For feeding, an essential neuronal structure in mammals connecting the body to the brain is the vagus nerve. Anatomical and functional studies have identified specific vagal ganglia that project to specific regions of the brainstem and segregate in a modality-specific manner (Bai et al., 2019; Borgmann et al., 2021; Gong et al., 2020; Grove et al., 2022; Prescott and Liberles, 2022; Ran et al., 2022; Ren et al., 2019; Williams et al., 2016; Zhao et al., 2022). Analysis of the SEZ, a center for processing sensory-motor information on taste and feeding related behaviors, is also being carried out in adult *Drosophila* using the strategy of marking cell types and following their projections to the brain (McKellar et al., 2020; Shiu et al., 2022; Sterne et al., 2021). These studies in mice and *Drosophila* however do not allow synaptic connections to be identified. In parallel, much progress has been made in identifying the different cell types in the brain at single cell level. For example, studies from various organisms and cellular contexts have shown that serotonin has wide-ranging effects on feeding, gut motility, mood and motor learning (Blundell, 1992; Cohen et al., 2015; Dayan and Huys, 2009; Gillette, 2006; Jacobs and Fornal, 1997; Okaty et al., 2019), and the complexity of the serotonergic neurons in the mouse brain is being characterized (Ogawa et al., 2014; Okaty et al., 2019; Pollak Dorocic et al., 2014; Ren et al., 2018). However, it is not known how the central serotonergic neurons are synaptically connected to specific sensory cells of the periphery.

Connectomic analysis with EM serial sections provides information on synaptic connectivity at different scales. In *Drosophila*, several volumes are being utilized, including adult hemi-brain, the whole brain, the ventral nerve cord; and the larval whole CNS volume (Ohyama et al., 2015; Phelps et al., 2021; Scheffer et al., 2020; Zheng et al., 2018). We have used the latter volume to map out the central circuits underlying feeding behavior, and to reconstruct the SEZ, a structure with analogy to the vertebrate brainstem and where the major sensory and motor neurons project to (Miroschnikow et al., 2018; Schlegel et al., 2016). A complete sensory and motor map has been reconstructed, including the topographical distribution of the sensory modalities from different regions, including the external, somatosensory, pharyngeal and enteric neurons. Furthermore, a group of serotonergic neurons in the SEZ that project to the enteric nervous system and the gut have been reconstructed in the CNS, and pointed out the striking similarities of the nerve that house these Se0 neurons to the mammalian vagus nerve (Schoofs et al., 2014b). However, despite the level and comprehensiveness of the connectome analysis in the fly feeding system, there is a major gap that prevents the next level of understanding even with an existing complete synaptic map of a brain region: one cannot identify the neurons at single cell resolution with their peripheral targets, for neither the sensory nor motor neurons. This knowledge would greatly advance understanding the brain map, since we can assign a biologically meaningful organ to which these projections belong. This would apply to mammalian systems as well, where synaptic mapping is being carried out for different brain regions in the mouse for example (Abbott et al., 2020). A whole animal EM volume, which includes the peripheral organs as well as the CNS, would fill this gap.

Here, we have used such a whole animal volume in *Drosophila* larva to assign all sensories from the peripheral feeding system to the brain at single cell and projection level. Through this, we elucidated the neuronal circuit for swallowing. The esophageal motor neurons driving this behavior are localized in the enteric ganglia, and passage of food along the esophagus is detected by Piezo positive mechanosensory neurons. The target of these mechanosensory neurons is a Se0 cluster of serotonergic neurons that in turn facilitates the activity of esophageal motor neurons, thereby stabilizing swallowing movements. There is in addition a direct synaptic connection from the esophageal motor neurons to the esophageal sensory neurons, revealing a neural substrate for an efference copy at synaptic level. Our results elucidate a circuit for swallowing that utilizes serotonin to stabilize a motor circuit in response to a successful and biologically relevant action. These also reaffirm the striking similarities between the circuit organization of mammalian and *Drosophila* vagus nerve (Schoofs et al., 2014b), a term used previously to describe the meandering nerve that connects the CNS with the periphery in insects (Newport, 1834).

## RESULTS

### STEM reconstruction of the *Drosophila* vagus nerve and enteric nervous system

To elucidate the synaptic map of the enteric nervous system in the brain, we used a whole animal STEM (scanning transmission electron microscopy) dataset that enables the reconstruction of all tissues of a *Drosophila* larva (**Figure 1A**). We fully reconstructed all neurons that connect the enteric nervous system (ENS) with the brain, which we refer to collectively as the vagus nerve (VN) of insects (Schoofs et al., 2014b) **(Figure 1B)**, a term used earlier to describe a nerve that projects from the brain to the gut in the moth *Sphinx ligustri* (Newport, 1834). The larval vagus nerve consists mostly of sensory projections (35) whose somas are located in distinct ganglia that comprise the ENS. All sensory projections target a special region of the SEZ, which we term the larval vagal center (_L_VC). The esophageal ganglia (EG) comprises 15 sensory neurons whose dendrites innervate the esophagus, while the hypocerebral ganglia (HCG) comprises five motor neurons that innervate the esophageal ring musculature (ERM) as well as eight sensory neurons that have tissue-unassociated local dendrites. The posteriorly most situated proventricular ganglia (PVG) connects the midgut and metabolic organs, including the gastric caeca and the nephrocyte-like garland cells (Weavers et al., 2009), via nine sensory neurons to the brain. In addition, nine neuromodulatory neurons in the PVG establish a connection between the midgut and the ring gland. Intriguingly, we also identified a novel sensory organ (Aorta_sens_) that innervates the aorta and sends projections to the SEZ and the median neurosecretory cells (**Figure S1**).

**Figure 1.**
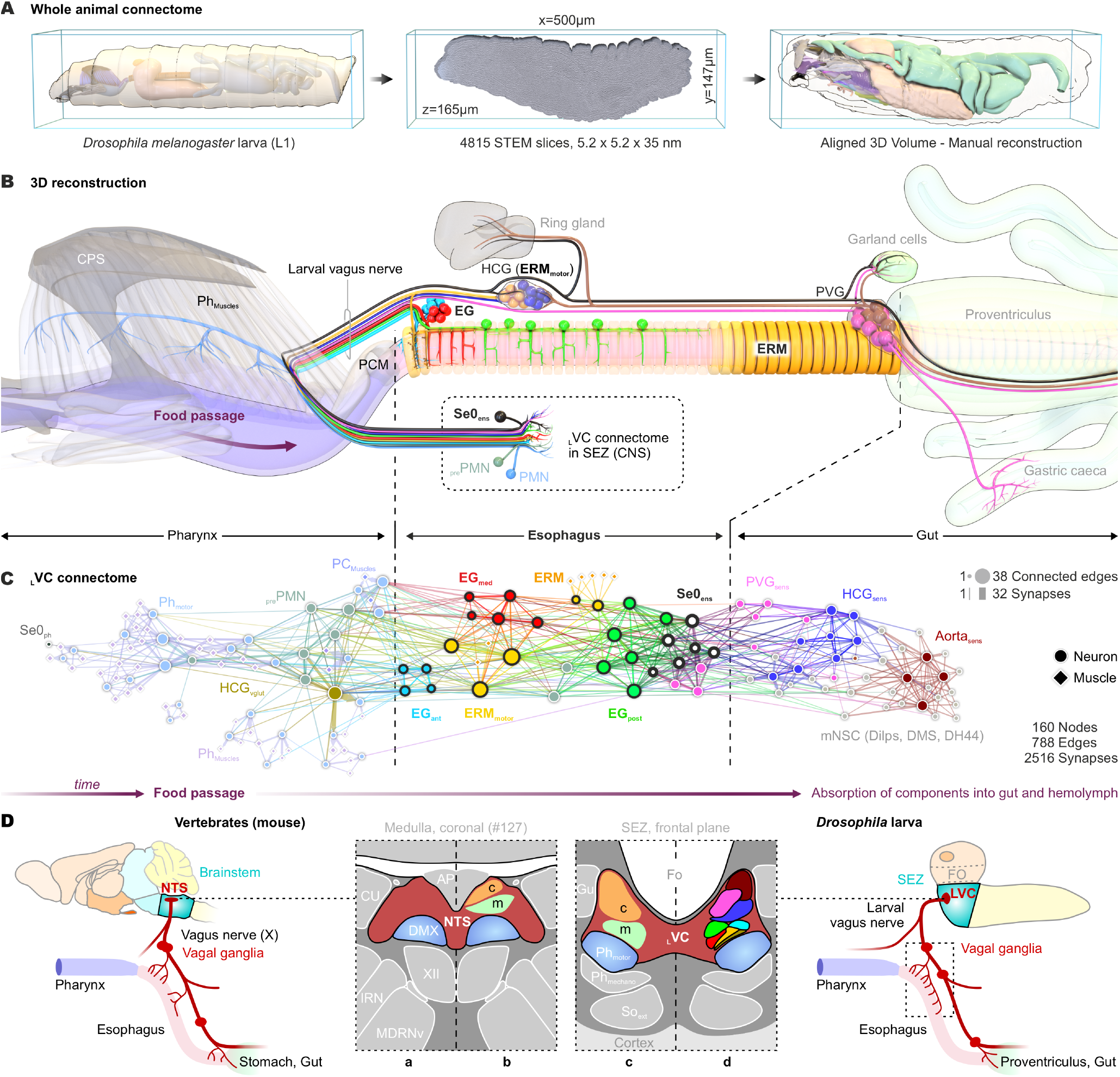
STEM-reconstruction of the larval vagus nerve and the enteric nervous system. **(A)** Whole first instar larva STEM (scanning transmission electron microscopy)-volume and manual reconstruction. **(B)** Three-dimensional reconstruction of the larval digestive tract and ENS which is interconnected with larval vagus center (_L_VC) in the SEZ (CNS) by the larval vagus nerve (Schoofs et al., 2014b; we refer here to the axonal pathway of CNS-ENS axis which projects through the antennal/recurrent nerve route as the larval vagus nerve). Reconstruction includes musculature and neuronal network for food ingestion and passage into the midgut. **(C)** Force-directed atlas of the enteric connectome in the SEZ correlates with coordinated actions of larval food intake behavior. **(D)** Comparison of the central representation of vagal afferents in mouse and *Drosophila* larva. In both species vagal mechano-(m) and chemosensory (c) inputs project to adjacent but distinct subareas in brainstem (mouse) and SEZ (*Drosophila*). Based on EM reconstruction the projection field of these afferents could be ascribed to specific groups of enteric sensory neurons in *Drosophila*. **Abbr**.: AP – area postrema, Aorta_sens_ – sensory neurons of the aorta, CPS – cephalopharyngeal skeleton, CU – cuneate nucleus, DH44 - diuretic hormone 44, Dilps - *Drosophila* insulin-like peptides, DMS - drosomyosuppresin, DMX – dorsal motor nucleus of the vagus nerve, EG_ant/med/post_ – esophageal ganglion (anterior, medial posterior), ENS – enteric nervous system, ERM – esophageal ring musculature, ERM_motor_ – esophageal ring musculature motor neurons, Fo - foramen, Gu – gustatory afferents, HCG – hypocerebral ganglion, IRN – intermediate reticular nucleus, MDRNv – ventral medullary reticular nucleus, mNSC – medial neurosecretory cell, NTS – nucleus of the solitary tract, PCM – pharyngeal constrictor musculature, Ph_muscles_ – pharyngeal musculature, PMN – pharyngeal motor neurons, pre – presynaptic site, _pre_PMN – pharyngeal premotor neurons, PVG – proventricular ganglion, Se0_ens_ – enteric Se0 neurons, So_ph_ – pharyngeal somatosensory afferents, So_ext_ – external somatosensory afference, SEZ – subesophageal zone, VN_ent_ – enteric vagus nerve, _L_VC - larval vagus center, XII – hypoglossal nucleus.

A distinct feature of *Drosophila* is a group of serotonergic neurons (Se0) located in the SEZ, a fly brainstem analog, that projects to all enteric ganglia, the midgut as well as the pharynx and ring gland, and has a role in foregut movements (Miroschnikow et al., 2018; Schoofs et al., 2014b). These serotonergic modulatory neurons are the only efferent connection of the CNS to the ENS and represent to date the only known source for peripheral serotonin in *Drosophila*.

We identified all monosynaptic connections between the enteric sensory and feeding related output neurons (motor, modulatory and neurosecretory), thereby defining the _L_VC connectome of the *Drosophila* larva. Force-directed graph mapping (**Figure 1C**) showed a striking concatenated series of connectivity modules interlinked along the foregut axis that mirror the temporal flow of food passage and the different phases of larval deglutition (Zinke et al., 1999), which can be divided into an oral, pharyngeal and esophageal phase similar to the mammalian swallowing system (Lang, 2009). The esophageal phase occupies a central position since it is the gateway for the most commonly used irreversible entry of food from the pharynx into midgut. The oral and pharyngeal modules are composed of pharyngeal premotor neurons and motor neurons in the SEZ that innervate different types of pharyngeal muscle. The esophageal phase is perceived by sensory neurons of the EG, motor neurons innervating the esophageal musculature and serotonergic modulatory neurons. The sensory neurons of the PVG, HCG, and the Aorta_sens_ form the last stage, namely the post-digestive phase of feeding and nutrient physiology.

Analysis of the mammalian vagus nerve projections in the brainstem provides a framework for comparing the organization of the _L_VC in the SEZ (**Figure 1D**). Classical anatomical work has defined topographically separated vagal sensory-motor regions in the brainstem, such as the nucleus of the solitary tract (NTS) and the dorsal motor nucleus of the vagus nerve (DMX). Through elegant molecular and functional analysis, the NTS could be further subdivided into topographically distinct target regions based on sensory modality and organ type (Williams et al., 2016), such as the separation of vagal mechanosensory and chemosensory projections from gastrointestinal tract via the enteric nodose ganglion. Previous connectomic work in *Drosophila* using the whole CNS volume showed that the _L_VC also displays a topographic separation (**Figure 1D**) (Miroschnikow et al., 2018). Using the current whole animal volume, the _L_VC targets in the brain could be elucidated at synaptic level and single cell resolution with respect to peripheral organs and ganglia (for details see **Figure S1B**).

### Mapping serotonergic output sites in the enteric nervous system

As a seed for functionally analyzing neuronal circuits underlying brain-gut interactions, we focused on the serotonergic modulatory Se0 neurons in the SEZ that innervate the entire ENS and have been shown to be involved in foregut peristalsis (**Figure 2A**) (Schoofs et al., 2014b). The EM reconstruction of the Se0 neurons revealed peripheral presynaptic sites with accumulation of clear core vesicles that lack distinct postsynaptic sites through the entire ENS (**Figure 2B; Figure S2A**). To analyze the primary release sites of serotonin in the ENS, we annotated each individual presynaptic site of the Se0 neurons and assigned them with a position tag relative to all enteric ganglia or innervation targets of the digestive system (**Figure 2C**). The analysis of the geodesic distance of the serotonergic release sites showed that the primary peripheral target of the Se0 neurons is the hypocerebral ganglion (HCG) and the proventricular ganglion (PVG) (**Figure 2D**). Expression analysis of all five serotonin receptors using knock-in T2A-Gal4 driver lines (Kondo et al., 2020) showed that serotonin receptors are widely distributed throughout the ENS and gut-associated peripheral organs (**Figure 2E**; Figures S2B-G and **Tables S1,S2**). Different subtypes are expressed in specific subsets in the sensory and motor neurons, as well as in muscles and gland cells (e.g., ring gland). To what extent serotonin becomes released throughout the projections, or specifically within these presynaptic structures is not known, but both are expected to act through volumetric transmission via serotonin receptor signaling.

**Figure 2.**
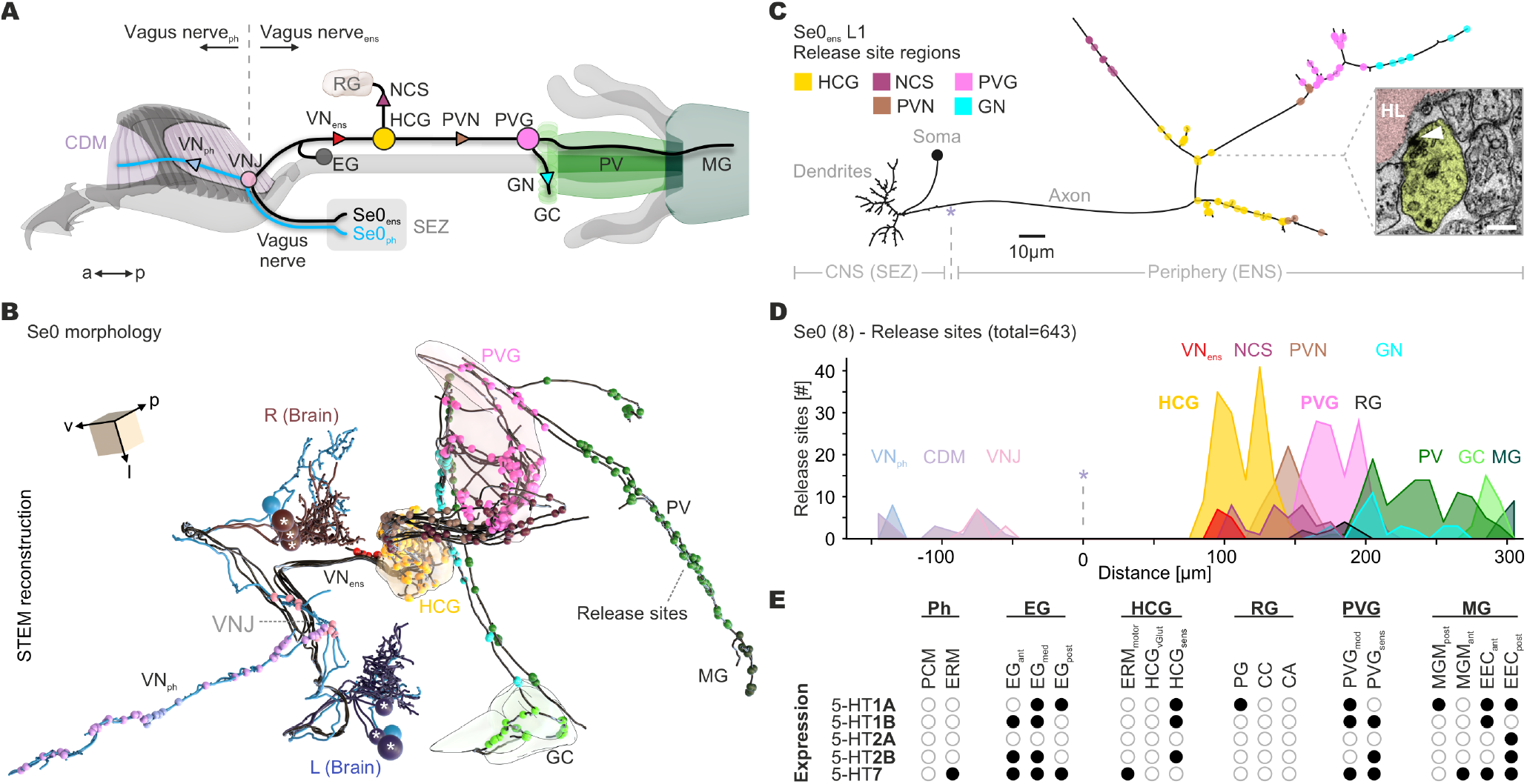
Serotonin release site and receptor subtype localization in the enteric nervous system. **(A)** Illustration of digestive tract and ENS marking the targets of Se0_ens_ and Se0_ph_ release sites. **(B)** STEM reconstruction of Se0_ens_ (asterisk) and Se0_ph_ and their axonic projections through the ENS. Colored spheres indicate region specific release sites. **(C)** Two-dimensional dendrogram of Se0_ens_ neuron (L1), colored circles indicate serotonergic release sites. Inserted EM-image shows a presynaptic site of a Se0_ens_ neuron without postsynaptic partners (scale bar: 0.2 μm). **(D)** Histogram shows spatial distribution of serotonergic release sites of all Se0 neurons onto digestive tract, ENS and endocrine organs. **(E)** Compendious serotonin receptor expression analysis of ENS (**Figure S2-S6 and Table S1**,**S2**). **Abbr**.: CA – corpora allata, CC – corpora cardiaca, CDM – cibarial dilator musculature, EG_ant/med/post_ – esophageal ganglion (anterior, medial, posterior), ENS – enteric nervous system, ERM – esophageal ring musculature, ERM_motor_ – ERM motor neuron, GC – garland cells, GN – garland nerve, HCG_vGlut/sens_ – hypocerebral ganglion (motor, VGlut, sensory), HL - hemolymph, MGM – midgut musculature, NCS – nervus cardio stomatogastricus, PCM – pharyngeal constrictor musculature, PG – prothoracic gland, Ph – pharynx, PVG_mod/sens_ – proventricular ganglion (modulatory, sensory), PVN – proventricular nerve, RG – ring gland, SEZ – subesophageal zone, VN_ent_ – enteric vagus nerve, VN_ph_ pharyngeal vagus nerve, VNJ – vagus nerve junction.

### Enteric motor neurons controlling esophageal peristalsis are modulated by serotonin

One of the serotonin receptors, 5-HT7, is expressed in motor neurons in the HCG that innervate the anterior ERMs. These motor neurons have a striking morphology and projection pattern: from their soma in the enteric ganglia HCG, one projection innervates the esophagus while the other projects to the brain in a bilateral fashion (**Figure 3A**). Their neurites in the SEZ have nearly equal amounts of input and output synapses, which places them as intermediates between other ENS neurons that can be clearly placed into input or output categories (**Figure 3B**). This suggests that the central projections of esophageal ring muscle motor neurons (ERM_motor_) relay information about the status of the esophageal musculature onto the neural network for feeding, in addition to acting through neuromuscular junction (NMJ) to control ERM activity **(Figure 3C**). A Janelia Gal4 line (GMR30F10) was identified which drives expression in the ERM_motor_ (**Figure 3D; Figure S3A**). These neurons are glutamatergic, as shown (1) by expression analysis of 5-HT7 in combination with Gal80 under the control of the VGlut promotor and (2) by VGlut fluorescent immunostaining (**Figures 3D,E**). STEM data further revealed that only the anterior seven ERMs are innervated by motor neurons, and the number of NMJs per ERM decreases from anterior to posterior (**Figure 3F)**.

**Figure 3.**
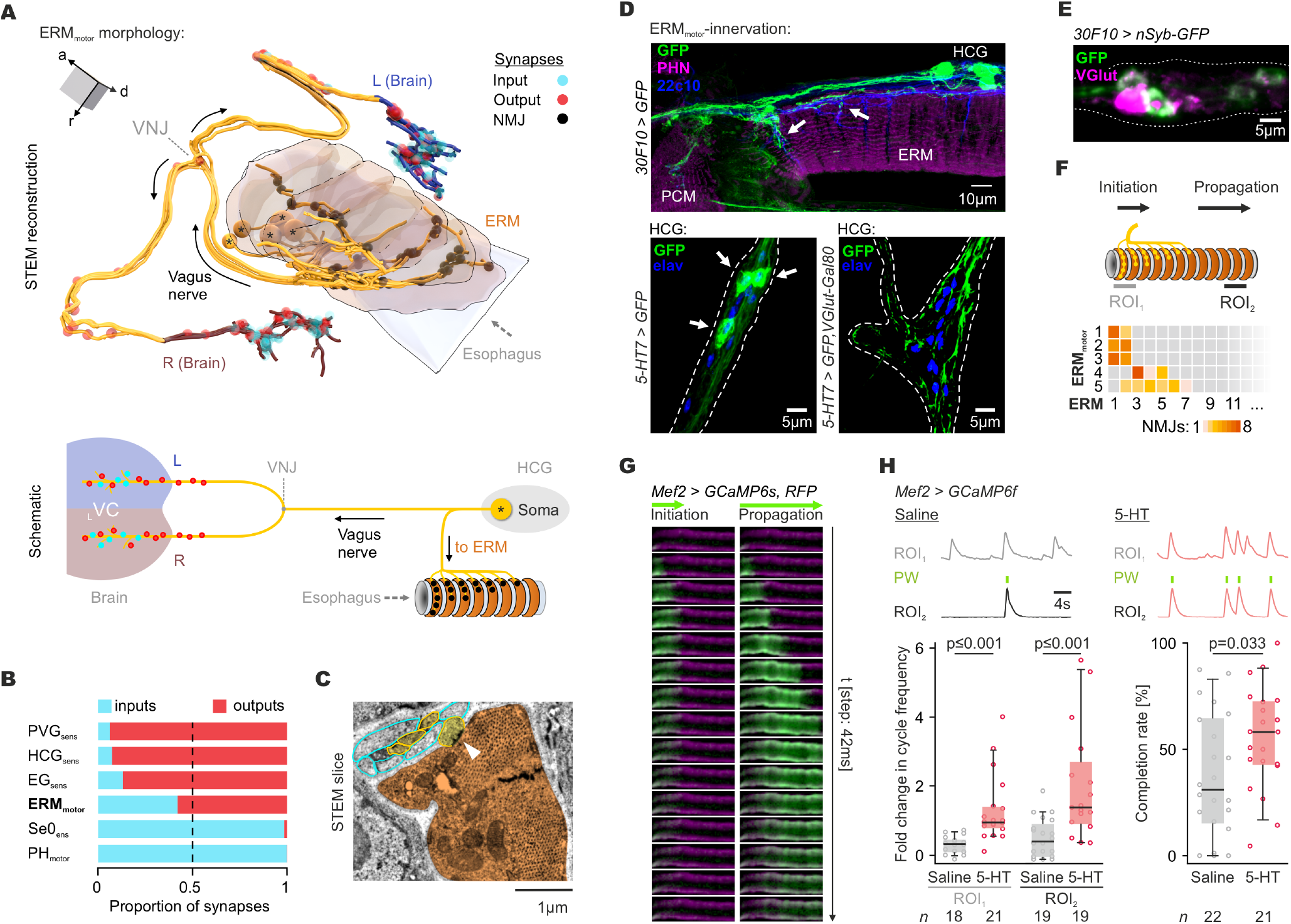
Effect of serotonin on esophageal ring muscle (ERM) motor system. **(A)** Top: STEM reconstruction of the motor neurons in the HCG showing the NMJs on the ERMs and motor afference to the CNS. Bottom: schematic drawing of ERM_motor_. **(B)** Bar plot shows the relation of input to output synapses in the CNS for sensory (input) neurons and modulatory/motor (output) neurons. Note the hybrid input to output proportion of ERM_motor_. **(C)** STEM slice of a motor axon (yellow) with NMJ (arrowhead) on a striated ERM muscle (orange). **(D)** Top: GFP-expression of *30F10-Gal4* covers ERM_motor_ in HCG. Bottom (left): expression of 5-HT7 showing three neurons in HCG. Bottom (right): expression of 5-HT7 in presence of Gal80 controlled by VGlut promoter shows three ERM_motor_ with 5-HT7. **(E)** Staining of 30F10 > nSyb-GFP shows colocalization with VGlut antibody. One VGlut-positive neuron shows no colocalization which is the VGlut positive HCG neuron (HCG_vGlut)_. **(F)** Top: asymmetric innervation of ERMs, number of NMJs per ERM. Bottom: heat map shows the number of NMJs per ERM for each ERM_motor_. NMJs are restricted to the initiation zone. **(G)** Calcium-imaging of ERMs revealed two distinct active muscle zones: initiation and propagation zones. **(H)** Top: calcium-recording of ROI1 (initiation) and 2 (propagation) showing the effect of serotonin on ERM-activity. Green lines mark complete peristaltic waves. Bottom: analysis showed an increase in ERM cycle frequency at ROI1 and ROI2 after application of serotonin (left). Analysis showed an increase in completion rate after application of serotonin (right). **Abbr**.: HCG - hypocerebral ganglion, ERM – esophageal ring musculature, ERM_motor_ – ERM motor neuron, _L_VC – larval vagus center, L - left, NMJ - neuromuscular junction, PCM - pharyngeal constrictor musculature, PHN - phalloidin, PW - peristaltic wave, R – right, ROI – region of interest, VNJ – vagus nerve junction.

We next monitored the activity of the ERMs using calcium imaging and observed two phases of esophageal peristalsis, which we termed “initiation” and “propagation” (**Figure 3G)**. Initiation phase can occur in the absence of propagation phase, and several rounds of initiation can occur before propagation proceeds; in contrast, we have never observed propagation in the absence of initiation. The positions of the two phases along the esophagus correspond with the neuroanatomical data on the innervation patterns, where the initiation phase coincides with the innervated region; the propagation phase is likely myogenic. Previous experiments indicated that activation of serotonergic Se0_ens_ neurons which is a Se0 subcluster projecting only to the ENS (**Figure S3C**) accelerates esophageal peristalsis by the release of serotonin (Schoofs et al., 2014b) (**Figure S3B**,**D**), and targeted knock down of serotonin synthesis by *TrhnRNAi* in Se0_ens_ neurons decelerates esophageal motility (**Figure S3E**). We carried out a series of ex-vivo experiments in which serotonergic activity was altered by exogenously added serotonin. Calcium imaging of the ERMs showed that both peristaltic phases (initiation and propagation) increased cycle frequency after application of serotonin (**Figure 3H, left panel**). Serotonin also increased the completion rate, which reflects the frequency with which a contraction wave progresses from initiation to propagation, resulting in a complete esophageal peristalsis (**Figure 3H, right panel**). Summarizing the findings for ERM_motor_ indicate that esophageal peristalsis is modulated by serotonin signaling from Se0_ens_ neurons through the enteric glutamatergic motor neurons via the receptor 5-HT7.

Next, we aimed to investigate the role of the central projections of ERM_motor_ based on their intermediate ratio of input and output synapses by performing lesion experiments with functional assays. Therefore, we first activated these neurons optogenetically at their somas using Chrimson, while observing esophageal muscle contractions (deglutition rate) and recording extracellularly from the larval VN (**Figure 4A**).

**Figure 4.**
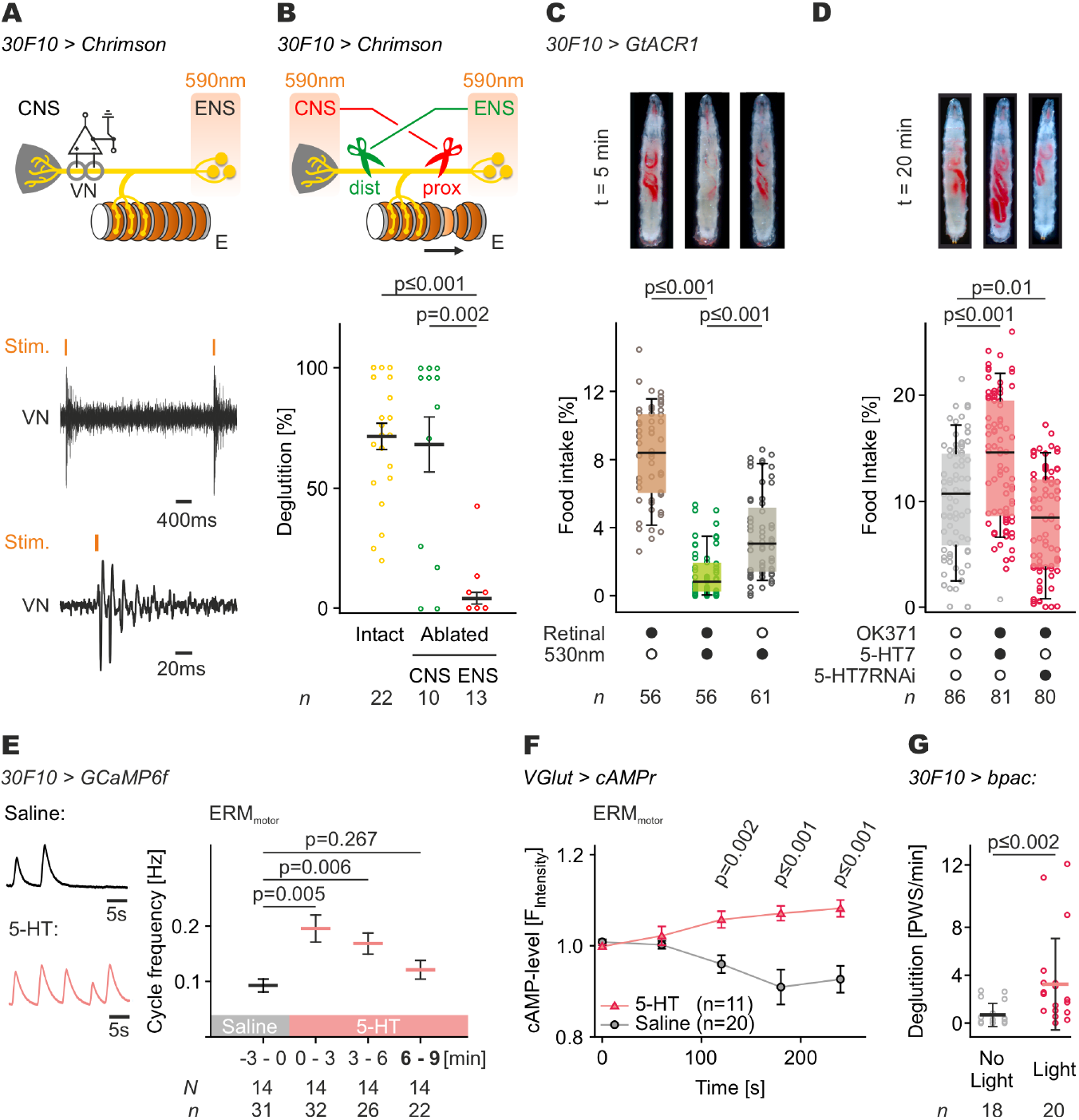
Effect of serotonin on esophageal ring muscle (ERM) motor system. **(A)** Activation of ERM_motor_ in ENS by Chrimson elicits afferent spikes detectable in VN-recordings. **(B)** Ablating the central neurite (dist) and activating ERM_motor_ in ENS by induced peristalsis. Ablation of primary neurite (prox) and activation in CNS abolished peristalsis. **(C)** Inhibition of the ERM_motor_ by GtACR1 suppresses food swallowing. **(D)** 5-HT7 overexpression in MNs by *OK371-Gal4* increased food intake, whereas the knock down of 5-HT7 decreased food intake. **(E)** Left: Calcium-imaging of ERM_motor_: representative data showing the effect of serotonin onto ERM_motor_. Right: analysis showed increased neuronal activity in ERM_motor_ after application of serotonin. **(F)** cAMP-reporter (cAMPr) showed an increased cAMP-level in ERM_motor_ after serotonin treatment. **(G)** Optogenetic increase of cAMP-level in ERM_motor_ by *UAS-bpac* accelerated deglutition. Data in (B), (F) show mean ± SEM and (G) shows mean + STD. Performed significance test: Mann-Whitney rank sum test. **Abbr**.: cAMP – cyclic adenosine monophosphate, dist – distal ablation (central neurite), ENS enteric nervous system, ERM – esophageal ring musculature, ERM_motor_ - ERM motor neuron, NMJ -neuromuscular junction, prox – proximal ablation (primary neurite), PW - peristaltic wave, ROI – region of interest, Se0_ens_ - enteric Se0 neurons, Se0_ph_ - pharyngeal Se0 neurons, VN – vagus nerve.

Activating ERM_motor_ elicited an afferent signal in the VN, indicating a functional connection to the CNS. To see whether the connection to the CNS was required for deglutition, we lesioned the distal part of the projection while leaving the connection to the esophagus intact (**Figure 4B**; green line). Lesion to the CNS at this point did not alter the deglutition rate, indicating that the CNS projection is not required for activation of ERM_motor_. By contrast, lesioning proximal to the junction site (while activating the CNS region) completely eliminated deglutition (**Figure 4B**; red line). Together, these findings indicate that for ERM_motor_, only the connection to the muscles is required for production of peristaltic waves, and that the connection to the CNS serves for afferent signaling of esophageal motor activity. Optogenetic inhibition of the ERM_motor_ neurons using GtACR1 led to a significant reduction of food intake (**Figure 4C**), further strengthening their functional role in regulating esophageal peristalsis.

To verify that the ERM_motor_ are modulated by serotonergic signaling via 5-HT7 (as we have already observed with the esophageal muscles), we genetically manipulated the activity of serotonergic receptor expression in these neurons. Overexpression of 5-HT7 in all glutamatergic neurons (via *OK371* driver) increased food intake; conversely, a knockdown of 5-HT7 expression decreased food intake (**Figure 4D**). We next carried out a series of ex-vivo analysis in which serotonergic activity was variously altered in these motor neurons. First, exogenously added serotonin increased the neural activity of the ERM_motor_ (**Figure 4E**) in the first three minutes, resulting in an increased number of peristaltic waves. The excitatory 5-HT7 is the only serotonin receptor which increases the intracellular cAMP level by activating adenylate cyclase (Witz et al., 1990). Therefore, we applied serotonin onto semi-intact larva expressing a cAMP sensor (cAMPr) in MNs and monitored the cAMP level of the ERM_motor_ **(Figure 4F)**. We observed an increase in the intracellular cAMP level of the ERM_motor_; similar results were obtained using a different cAMP sensor (Epac-camps; **Figure S4**), strengthening the view that serotonin modulates ERM_motor_. We also optogenetically modulated the cellular cAMP level in the ERM_motor_ by using the photoactivatable adenylate cyclase, bPAC (Stierl et al., 2011). Monitoring the esophageal peristalsis showed that increasing cAMP activity in the ERM_motor_ caused increased peristaltic movements **(Figure 4G)**. Taken together, these data show that esophagus peristalsis is driven by ERM_motor_ whose activity can be modulated by serotonin via 5-HT7.

### Distinct classes of esophageal Piezo mechanosensory neurons sense food passage through the foregut

Having identified the esophageal motor neurons of the swallowing motor system and demonstrated their modulation by serotonin, we next wanted to determine which sensory neurons play a role in esophageal peristalsis. We therefore identified in the whole animal STEM volume all sensory neurons with dendritic contact to the esophagus which are located in the EG (**Figure 5A)**. Subsequent clustering analysis by neuronal morphology and synaptic connectivity of the total enteric connectome, including the pharyngeal motor neurons (PMNs), revealed three distinct clusters of neurons in the EG along the anterior-posterior axis: the EG_ant_, EG_med_ and EG_post_ (**Figure S5A**). Notably, the positions of the different sensory neurons along the esophagus correlate with the previously described dynamics of esophageal muscle activity (**Figure 3G**), namely EG_ant_ and EG_med_ in the initiation zone and EG_post_ in the propagation zone. The examination of the synaptic connectivity between all neurons of the EG showed that the intra-cluster connection clearly predominates over the inter-cluster connections, confirming that the three esophageal clusters represent independent sensory units (**Figure S5B**). Since the three distinct esophageal units show strong reciprocal synaptic intraconnections, this may imply that upon perception, neural response of one neuron conditions the activity of all neurons within a unit, resulting in an amplification or synchronization of sensory inputs to downstream targets. The analysis of the EG connectivity also revealed that directed synaptic connections between EG neurons occur only to a minor extent, indicative of a non-hierarchical organization in the sensory perception.

**Figure 5.**
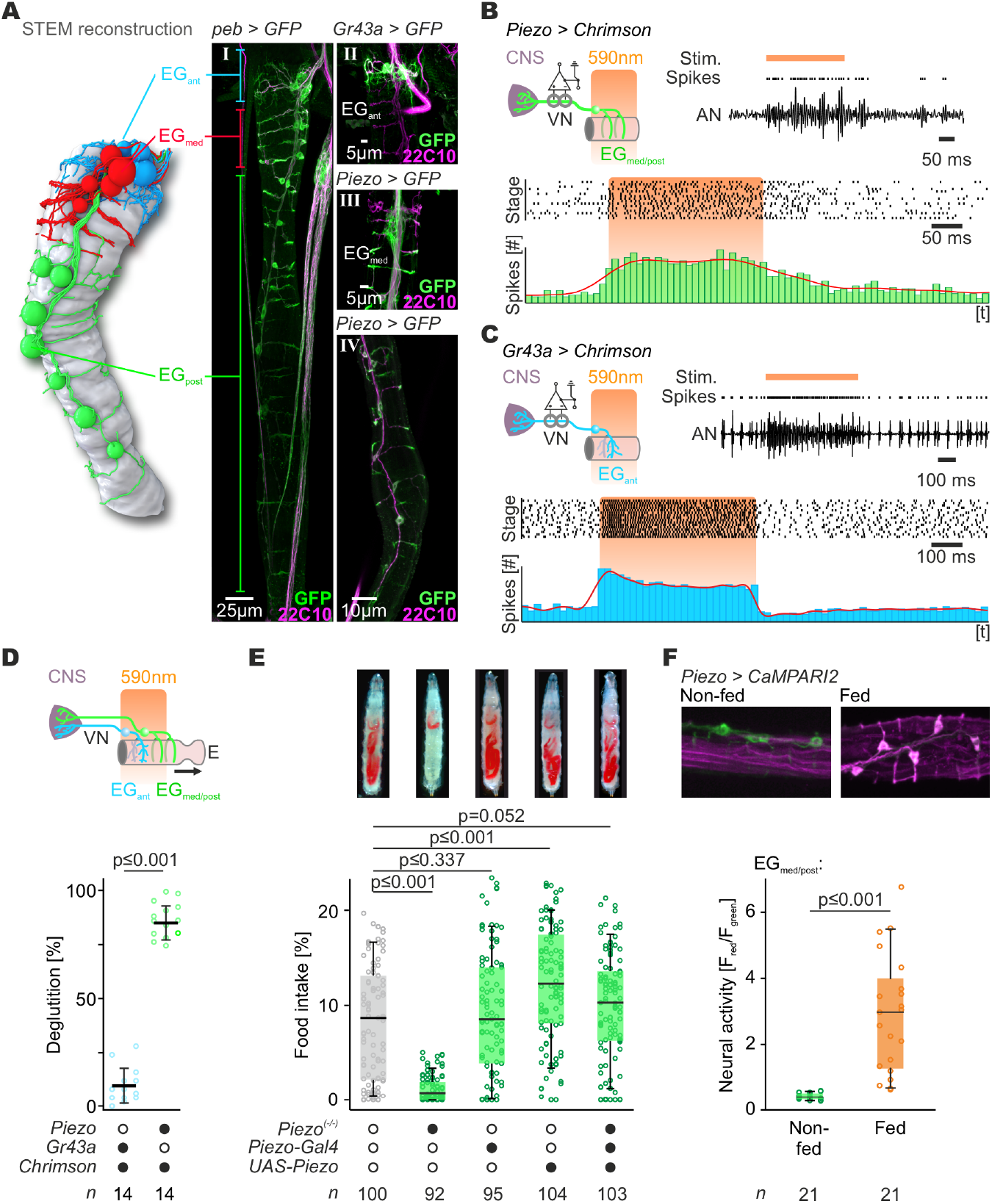
Piezo-expressing neurons in EG monitor food passage. Left: STEM reconstruction of EG. Right: GFP-staining of whole EG (I). Three different clusters of the ganglion are shown: EG_ant_, EG_med_ and EG_post_. Chemoreceptor Gr43a is expressed in EG_ant_ (II) whereas EG_med_ (III) and EG_post_ (IV) express the mechanoreceptor Piezo. **(B**,**C)** Activation of Piezo-expressing (**B**) and Gr43a-expressing (**C**) neurons in EG using Chrimson elicit afferent spikes to CNS. **(D)** Activation of Piezo-expressing EG neurons elicits peristalsis, whereas Gr43a-expressing EG neurons do not. Data are mean ± SD. **(E)** *Piezo*^*(-/-)*^ larvae showed reduced food intake which was rescued by expression of Piezo via Gal4/UAS-system in *Piezo*^*(-/-)*^ background. **(F)** CaMPARI-experiments of EG_post_ using *Piezo*^*(KI)*^*-Gal4* indicated that the mechanosensory EG_post_ neurons are capable of monitoring food passage through the esophagus. Performed significance test: Mann-Whitney rank sum test. **Abbr**.: EG_ant/med/post_ - esophageal ganglion (anterior, medial, posterior), Stim. - stimulus, VN – vagus nerve.

Marker stainings with receptors showed that the chemoreceptor Gr43a is expressed only in EG_ant_, while the mechanoreceptor Piezo is expressed in EG_med_ and EG_post_ (**Figure 5A**). The Gr43a expressing EG_ant_ neurons show to a minor degree direct monosynaptic connections to PMNs and premotor neurons (_Pre_PMNs) but not ERM_motor_. In contrast the Piezo expressing EG_med_ neurons have monosynaptic contacts with ERM_motor_, which may indicate a role in the initiation and coordination of esophageal peristalsis, because the receptive fields of EG_med_ lie posterior to the neuromuscular terminals of ERM MN 1-3 at the esophagus. The most intriguing EG is the Piezo-expressing EG_post_ which has monosynaptic connection to the modulatory Se0_ens_, and are well positioned to putatively detect the irreversible transport of food to the midgut.

To address the function of the esophageal sensory neurons with respect to swallowing motor pattern, we performed combined optogenetic and electrophysiological experiments. Activation of either Piezo or Gr43a positive EG neurons with Chrimson elicited spikes in the larval VN, indicating that an afferent sensory signal is transmitted to the CNS by both types of sensory neurons (**Figure 5B,C**). Critically, however, only the excitation of Piezo neurons by Chrimson, but not Gr43a neurons, induced deglutition (peristaltic waves) (**Figure 5D**), indicating that mechanosensory inputs have a stronger impact on the output system for esophagus motility as compared to chemosensory inputs. To determine whether *Piezo* gene activity is required in these mechanosensory neurons, we performed behavioral assays with *Piezo*^*(-/-)*^ animals. Indeed, these had significantly decreased food intake in behavioral assays (**Figure 5E**), showing the importance of these mechanosensory input for feeding behavior in vivo. Next, we addressed what these Piezo-expressing EG neurons are sensing using the calcium integrator CaMPARI. Strikingly, significant increase in activity of EG_post_ was seen only in conditions when animals are feeding (**Figure 5F; Figure S5C**). These anatomical and functional studies indicate that mechanosensory Piezo expressing EG neurons send information to the CNS concerning the presence of food in the esophagus. At the current stage of the whole larva STEM reconstruction, we cannot quantify the weight of the polysynaptic integration of these sensory neurons, but it is known that integration of chemosensory inputs, e.g. taste perception, occurs exclusively through interneurons in comparison to mechanosensory inputs (Miroschnikow et al., 2018).

### Influence of direct and indirect sensory pathways onto Se0_ens_

We next analyzed the monosynaptic connections of enteric sensory and other interoceptive neurons onto the three classes of feeding related output neurons in the brain: PMNs, serotonergic modulatory neurons and neurosecretory neurons (**Figure 6A,B**). This revealed a clear division of labor between the enteric ganglia and their targets. For output neurons, Se0_ens_ neurons receive their main sensory input from EG_post_ and PVG_sens_ neurons, the median neurosecretory cells (mNSCs) from HCG_sens_ and Aorta_sens_ neurons. In contrast, the pharyngeal system receives weak synaptic input from EG_ant/med_ and HCG_vGlut_ neurons. For input neurons, nearly all distinct sensory clusters receive their main synaptic input through axo-axonic intra-cluster connections. Strikingly, the Piezo-expressing EG_post_ neurons receive their primary input from ERM_motor_ neurons, representing an efference copy.

**Figure 6.**
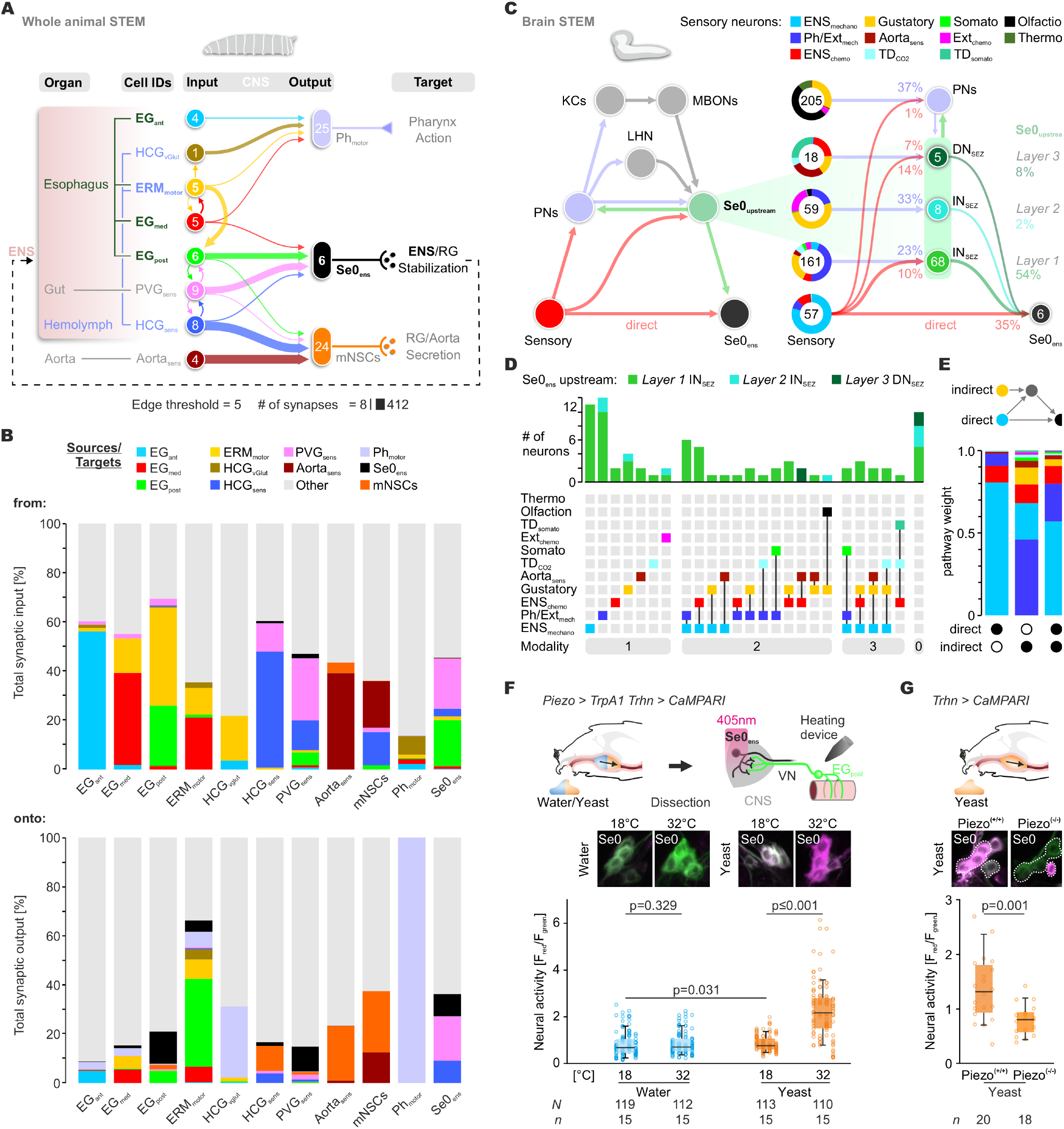
Direct and indirect sensory pathways onto the central serotonergic Se0_ens_ neurons. **(A)** Illustration of the monosynaptic enteric synaptic connections to the three primary feeding related neuronal output systems. **(B)** Color bar plots show the total synaptic input and output in percentage of different neuronal groups relevant in the swallowing circuit. **(C)** Illustration of all direct and indirect (via interneurons) sensory pathways onto the Se0_ens_ neurons. Numbers within circles represent the number of neurons. Percentage sensory composition (the five left donut circles) is shown by sensory modalities. **(D)** Graph represents number of Se0_ens_-upstream neurons and integrated sensory modality for the indirect (polysynaptic) integration of sensory inputs onto Se0_ens_ neurons by labeled lines and multimodal pathways. **(E)** Bar plots show the weight for all direct, indirect and combined direct/indirect sensory pathways onto the Se0_ens_ neurons. Colors represent the proportion of the different sensory modalities. **(F)** CaMPARI-experiments: local stimulation of Piezo-expressing EG_post_ neurons directly after feeding but not non-feeding phase resulted in significantly increased neural activity of Se0 neurons. Performed statistical test in **A/B**: Mann-Whitney rank sum test. **(G)** CaMPARI-experiments: Se0 neurons in *Piezo*^*(-/-)*^ background showed reduced neural activity while ingesting yeast.

The whole larval volume connectivity is based only on ENS-specific direct input-to-output connections. In order to identify all direct and indirect polysynaptic pathways from sensory neurons onto Se0_ens_ neurons, we utilized earlier EM studies on whole CNS volume to see how ENS-independent sensory information and higher brain regions, like mushroom body and lateral horn, are layered onto the basic ENS-Se0_ens_ connectome (**Figure 6C**) (Eichler et al., 2017; Hückesfeld et al., 2021; Miroschnikow et al., 2018; Winding et al., 2023). The polysynaptic integration of first-order sensory information comprises labeled-lines and multimodal pathways through second-order interneurons that connect directly to Se0_ens_. These indirect pathways include all different sensory modalities/sources, e.g. feeding related pharyngeal and external gustatory and mechano-sensory inputs (**Figure 6D**). To scale the contribution of the distinct inputs of the Se0_ens_ connectome, we calculated the normalized weight for each direct and indirect (only via Se0_ens_ upstream interneurons) sensory pathway (**Figure 6E**), showing that ENS_mechano_ and Ph/Ext_mechano_ are the predominant sensory inputs to the Se0ens, whereas ENS_chemo_ and gustatory inputs are integrated to a moderate extent.

To test for functional connection, we focused on the EG_post_ neurons that monitor food passage. We thus activated the EG_post_ with TrpA1 and monitored the activity of the Se0 neurons with CaMPARI. This was done in two different conditions, one with water (low feeding) and one with yeast (high feeding). When presented with water, activation of EG_post_ did not result in significant increase in Se0 activity; by contrast, performing these experiments with yeast resulted in significantly increased Se0 activity (**Figure 6F**). This is supported by results in which the neural activity of Se0 neurons was increased upon presenting attractive nutrients and suppressed upon aversive ones (**Figure S6A**). Furthermore, application of the same attractive nutrient with different viscosities indicates that the neural response of Se0 neurons is not solely due to gustatory inputs, but combined mechano-gustatory inputs. Experiments in which food experience was limited to gustatory perception showed a reduced neural response of Se0 neurons, further pointing to a considerable mechanosensory input onto the Se0 neurons (**Figure S6B-D**). Furthermore, in Piezo^(-/-)^ animals, the induction of Se0 activity upon yeast feeding is no longer observed (**Figure 6G**), demonstrating a critical role of Piezo-dependent activation of Se0 neurons. Summarizing the synaptic and functional data, the circuit architecture suggests a mechanism by which the Se0_ens_ neurons respond primarily to mechanosensory inputs from the EG_post_ neurons that food has successfully moved through the esophagus, but their activity is also dependent on food quality, e.g. taste and texture.

### Synaptic representation of afferent and efferent copy pathways in the serotonergic swallowing circuit

We next generated an information flow map with the EG neurons, ERM_motor_ neurons and the Se0 neurons. Asides from the intra-sensory and intra-motor connections, the circuit flow has two key elements. One is an axo-axonic efference copy whereby the ERM_motor_ neurons connect to the EG_post_ neurons; the second is an axo-dendritic afferent connection in which the EG_post_ neurons connect to the modulatory Se0 neurons (**Figure 7A**; see also Figure 6A,B). To delineate the structure of these connections in more detail, we considered the arrangement of synapses on the neurites as a key determinant for the neuronal response in the circuit (Rall, 1995). Thus, we calculated the geodesic distance for each synaptic input and output of each ERM_motor_ and EG_post_ neuron using the vagus nerve junction (VNJ) as collective origin. For the Se0_ens_ neurons, we used the CNS entry site of the VN as the origin for the measurement (**Figure 7B,C**). This analysis revealed that the axo-axonic connections from ERM_motor_ onto EG_post_ show the lowest geodesic distance compared to all other inputs and outputs, strengthening our assumption of an efference copy (**Figure 7C**). Finally, for Se0_ens_ neurons we identified that the axo-dendritic inputs of EG_post_, along with all other enteric ones, are closer to the anticipated locus of spike initiation (CNS entry site) than synaptic sites of all other neurons presynaptic to Se0_ens_ (**Figure 7B-D**). Therefore, we consider that the enteric inputs, in particular EG_post_, are likely to have a superordinate influence on the neural activity of Se0_ens_.

**Figure 7.**
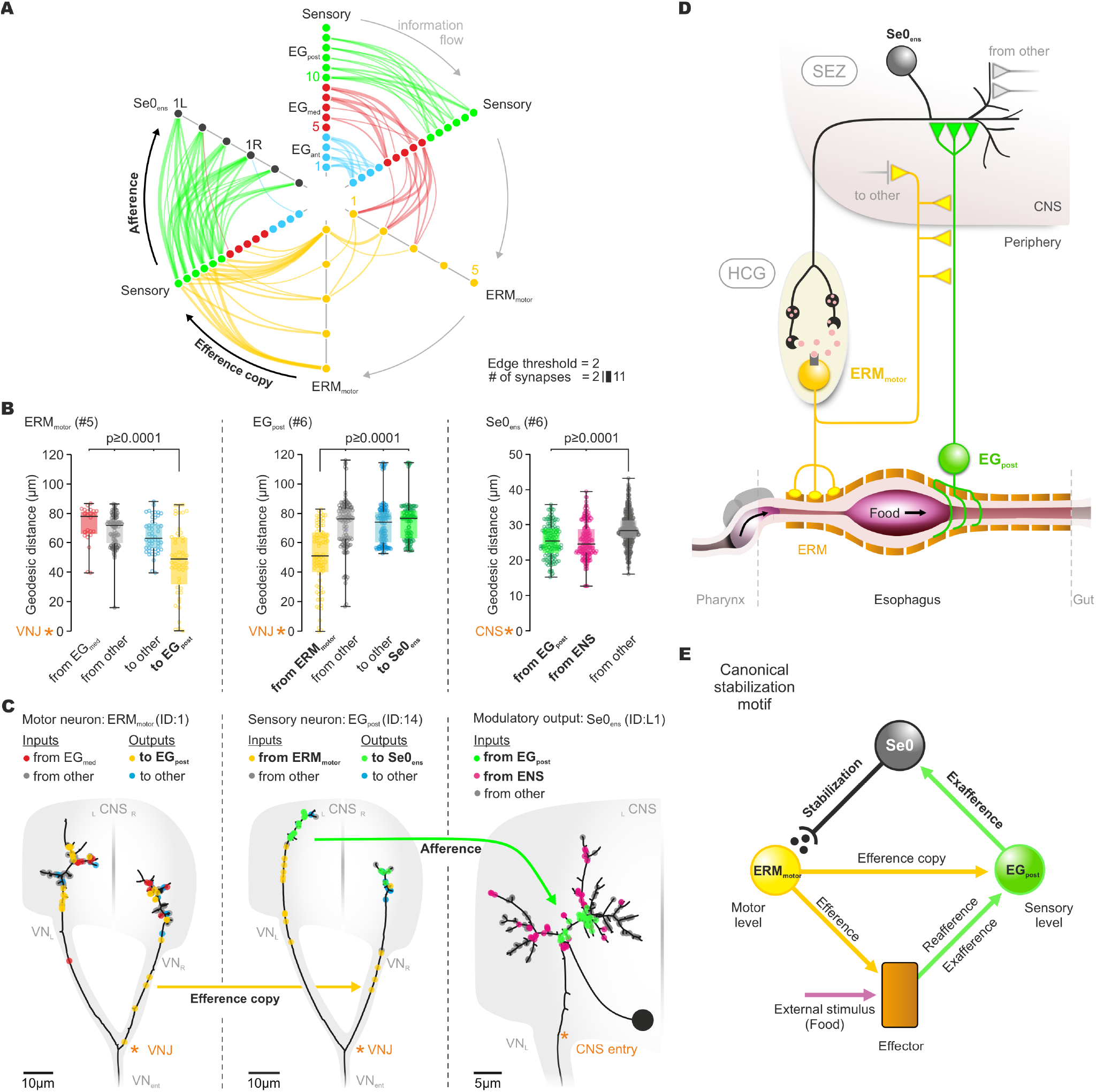
Reinforcement motif of the swallowing circuit. **(A)** Radial information flow diagram of sensorimotor circuit for swallowing. **(B)** Spatial synapse analysis for ERM_motor_, EG_post_ and Se0_ens_. Left: ERM_motor_ outputs to EG_post_ neurons are located in front of all other output- or input sites. Middle: EG_post_ input sites from ERM_motor_ are prior to outputs onto Se0_ens_ neurons. Right: inputs from EG_post_ neurons show significantly lower geodesic distance compared to other inputs. Performed statistical test: one-way ANOVA. **(C)** Two-dimensional dendrograms of sensorimotor circuit for swallowing: axo-axonic inputs (yellow) of ERM_motor_ (left) onto EG_post_ neurons (middle) resemble an efference copy. Following inputs (green) of EG_post_ neurons onto Se0_ens_ neurons (right) indicate a reinforced signaling, based on their spatial assembly to Se0_ens_ axo-dendritic transition zone (locus of spike initiation). **(D)** Circuit architecture for spatial sensory integration along the esophagus of EG_post_ neurons during food swallowing. Food passage is detected by mechanoreceptive EG_post_ neurons which reinforce Se0 neurons to release serotonin. ERM motor system is stabilized by released serotonin, and ERM motor output functions as efference copy (EC) for EG_post_ neurons. **(E)** Illustration of the canonical stabilization motif for the food swallowing motor program. **Abbr**.: EG_post_ – posterior esophageal ganglion, ERM – esophageal ring musculature, HCG – hypocerebral ganglion, VN – vagus nerve.

Taken together, our work elucidates an elemental circuit motif for reinforcement of the swallowing motor program by serotonin. At its core, the swallowing circuit consists of a motor neuron in the enteric ganglia that innervates the esophageal muscles, a mechanosensory neuron along the esophagus that detects passage of food, and a serotonergic neuron in the CNS that receives this sensory information and facilitates the motor neurons. Thus, this circuit enhances the swallowing movements when the nervous system registers that food has successfully passed into the body, and may represent an ancient form of motor reinforcement learning **(Figure 7E**).

## DISCUSSION

The seemingly simple act of swallowing is arguably the single most salient decision an animal has to make. The motor system underlying swallowing is the ultimate “final common path” (Sherrington, 1906) for feeding behavior, as it is the last and essentially irreversible step when food is taken into the body. We have elucidated the elemental neuronal circuit for swallowing in *Drosophila*, through the first use of a whole animal STEM volume that enabled the identification of all feeding relevant connections between the body and the brain. The circuit comprises motor and sensory neurons in the esophagus that are interconnected with central serotonergic modulatory neurons, and whose motor pattern becomes stabilized upon successful food swallowing.

The esophageal ring musculature motor neurons (ERM_motor_) underlying swallowing are located peripherally in an enteric ganglia and innervate the esophageal muscles along the anterior-posterior axis in a graded fashion. These pseudounipolar neurons have a striking morphology: in addition to innervating the muscles, the ERM_motor_ neurons have synaptic outputs in the brain, and to the posterior esophageal ganglia sensory neurons (EG_post_). This shows, in agreement with our functional experiments, that ERM_motor_ neurons can send an afferent motor activity signal back to the brain and the sensory system, thus providing a neuronal substrate of an efference copy at single-cell and synaptic level. The EG_med_ and EG_post_ neurons are arrayed in a sequential manner along the length of the esophagus and become activated during swallowing. These neurons express the Piezo mechanoreceptor, suggesting that they sense food passage through distension of the esophagus, and mutation in the *Piezo* gene eliminates their response to food. The differing monosynaptic connectivity of all EG sensory clusters allows a clear assignment of their functional roles in swallowing. The anteriorly located esophageal sensory neurons, EG_ant_ and EG_med_, show direct connections to the pharyngeal motor system and ERM_motor_ neurons, respectively; this points to a reafferent pathway that contributes to the coordination of the swallowing motor patterns. However, the mechanosensory EG_post_ neurons have a high number of synaptic outputs to a cluster of central serotonergic neurons in the CNS (Se0_ens_) that modulates esophageal peristalsis. This afferent signal onto the Se0_ens_ neurons likely represents a “quality control” of swallowing, which can be deduced from the synapse topology upon comparing the location of the axo-axonic ERM_motor_-to-EG_post_ to the axo-dendritic EG_post_-to-Se0_ens_ connections. Signal-integration of the more anteriorly located ERM_motor_-to-EG_post_ connections (efference copy) onto the mechanosensory response due to esophageal distension, potentially prevents the influence of sensory signals caused by self-initiated movements (“reafference”). After the subtraction of the efference copy, the residual mechanosensory signal of EG_post_ onto SE0_ens_ would be identified as “exafference” and perceived as being elicited by the passage of food through the esophagus. In other words, this elemental circuit architecture provides a single cell resolution neuronal substrate for how animals can distinguish its own self-generated action versus that from extrinsic stimuli, which is necessary to achieve a biologically relevant goal like feeding.

Optimal feeding is not solely based on a coordinated series of sensory-motor actions that respond to a single sensory signal at a given moment in time. Rather, the trajectory of an organism’s actions changes as the perception of the multisensory food signals become altered as feeding progresses. This is combined with processing through higher brain regions, such as from the lateral horn (innate behavior) and the mushroom body (associative memory), to create a meaningful and biologically relevant context. Brain-wide analysis of polysynaptic sensory-to-Se0_ens_ integration revealed that all second-order interneuron pathways to Se0_ens_ convey sensory information in the form of labeled lines or multimodal combinations. Some of these interneurons form convergence pathways for on-going as well as stored sensory information, e.g. from mushroom body output neurons (Eichler et al., 2017; Miroschnikow et al., 2018). Although both the mono- and polysynaptic sensory pathways integrate predominantly mechanosensory information of enteric and pharyngeal origin, our functional experiments indicated that mechanosensory inputs are essential but not sufficient to activate Se0_ens_ neurons. Not surprisingly, reinforcing signals onto the swallowing motor system only occur when an attractive nutrient has been ingested. The final common target of the swallowing system is the Se0_ens_ neurons, which in turn project to the entire enteric nervous system, the neuroendocrine organs and the midgut. A main functional target of Se0_ens_ neurons is the ERM_motor_ neurons, which express the excitatory serotonin receptor 5-HT7. Manipulating receptor signaling by genetic, ex-vivo and optogenetic methods indicate that serotonin facilitates ERM_motor_ neuron activity. Based on these data, we propose that the core swallowing system represents a fundamental circuit motif for reinforcing a biologically meaningful motor action, using serotonin as a reward molecule to stabilize the swallowing motor pattern.

Stabilizing motor activity has been proposed to be a fundamental organizing principle underlying serotonin function (Allman, 1999). The serotonergic architecture underlying the swallowing motor program could represent a blueprint around which additional circuit elements can be organized to reinforce a biologically meaningful action. Work of Jacobs and Fornal (1997) showed that neural activity of certain serotonergic neurons which are activated by somatosensory and proprioceptive stimulation is associated with motor output activity, suggesting a facilitating effect of serotonin on motor function. Serotonergic neurons are also involved in controlling many motor activities in *Drosophila*, including feeding related behaviors (Albin et al., 2015; Eriksson et al., 2017; Schoofs et al., 2014a; Yao and Scott, 2022). In *C. elegans*, enteric serotonergic neurons respond to food ingestion and modulate the feeding circuit (Rhoades et al., 2019; Song et al., 2013). In *Aplysia*, the sensitization of the gill withdrawal reflex has three basic components (a mechanosensory neuron, a motor neuron and a modulatory serotonergic neuron) whose coordinated activity is required for memory formation (Kandel, 2001; Upreti et al., 2019). In zebrafish, serotonergic neurons may be involved in switching between flexible actions, and focused prey (Kawashima et al., 2016; Marques et al., 2020). In song bird motor learning (Fee and Goldberg, 2011; Mackevicius and Fee, 2018), the circuit utilizes dopamine rather than serotonin but the essential features of the system can be discerned: an action generating circuit that connects to the sensory system via efference copy, which is coupled to a reward system to reinforce a vocal motor pattern. What has been lacking in almost all these cases is the identity of the sensory neurons that provide direct or indirect synaptic inputs to the serotonergic neurons within functional circuits, and how they are connected to the motor neurons at synaptic and single cell level. The serotonergic swallowing circuit architecture presented here could be used as a foundational building block for understanding how modulatory neurons can be integrated into functionally defined sensory-motor circuits to facilitate and stabilize rewarding behaviors.

Finally, as previously noted, the term “vagus nerve” was used nearly 200 years ago to describe a nerve in an insect that traverses from the brain down to the gut, with branching projections onto various parts of the peripheral organ (Newport, 1834). The similarities between insect and mammalian systems are remarkable. In both cases, a single nerve directly connects the gut with the brain. In mammals, the sensory organs are localized in distinct ganglia, with dendrites onto various peripheral organs involved in internal physiology; such organization is also found in *Drosophila*. The vagal sensory neurons project to the brainstem and segregate according to modality and organ origin; there is also segregation in the SEZ based on sensory modality and organ identity of the larval vagal sensory neurons. Despite the differences in the number and cell types, the core brain-body circuits involving serotonin and enteric sensory neurons to stabilize feeding related motor patterns may also be operating in mammals.

## ACKNOWLEDGEMENTS

We thank Bubu Hückesfeld, Nicole Kucharowski, Ekatarina Nikitina, Anne Oepen, and Fabian Zink for help with cloning and testing of constructs and fly lines. We thank Laurin Büld, Marek Eckart, and Jan Weis for help with behavior experiments. We thank Michael J. Texada, Hermann Aberle and Jan Veenstra for sharing fly lines, antibodies and resources. We especially thank Andreas Thum and Stephen Liberles for helpful comments on an earlier version of the manuscript.

## AUTHOR CONTRIBUTIONS

Investigation, A.S.;Transgenic fly generation, I.Z.;

EM-dataset: C.S.M., A.C.; EM-reconstruction: A.S., A.M.;

Computation/Coding: P.S.;

Analysis: A.S., A.M., P.S.;

Visualization: A.S., A.M.;

Supervision: M.J.P.;

Writing – original draft: A.S., M.J.P.

## DATA AND MATERIAL AVAILABILITY

All data generated or analyzed during this study are included in the manuscript and supporting files. We used the whole animal STEM volume reported in Pearl et al., (in prep.). Contact for dataset accession: A. Cardona.

## MATERIAL AND METHODS

### Fly work and fly lines

All larvae were kept on 25 °C under 12 h light/dark cycle if not otherwise stated. For behavioral experiments 4 h egg collections were made on apple juice agar plates containing a load of yeast/water paste. After 48 h, larvae were transferred into vials (60 larvae per vial) containing standard cornmeal medium. For other experiments, e.g. functional imaging, electrophysiological recording and antibody staining, 4 h egg collection were made in vials with standard cornmeal medium with a spot of yeast/water paste and afterwards kept for four days on 25 °C. Only larvae for optogenetic stimulation were raised on fly food containing 150 μM all-trans retinal and kept under dark conditions. All larvae used for the experiments were 96±2 h old. The following *Drosophila melanogaster* lines were used (see also Table S3 for genotypes of experimental flies):

#### Driver lines

*5-HT1A*^*2A-Gal4*^, *5-HT1B*^*2A-Gal4*^, *5-HT2A*^*2A-Gal4*^, *5-HT2B*^*2A-Gal4*^, *5-HT7*^*2A-Gal4*^ (all (Park et al., 2018) and (Kondo et al., 2020)), *30F10-Gal4* (BDSC #49643), *Gr43a*^*Gal4*^ (BDSC #93447), *Mef2-Gal4* (BDSC #27390), *OK371-Gal4* (BDSC #26160), *peb-Gal4* (BDSC #80570), *Piezo-Gal4*^*IIA*^ (BDSC #58771), *Piezo-Gal4*^*III*^ (BDSC #59266), *Piezo*^*Gal4*.*KI*^ (BDSC #78335), *Se0*_*ens*_*-Gal4* (BDSC #47343), *Se0*_*ph*_*-Gal4* (named “*mn9*” in (McKellar et al., 2020)), *Trhn-Gal4* (BDSC #38389), *Trhn-lexA (Alekseyenko et al., 2014), VGlut-GAL4* (BDSC #24635).

#### Effector/reporter lines

*lexAop-CaMPARI* (for generation see below), *UAS-5-HT7* (Kerr et al., 2004), *UAS-5-HT7-RNAi* (BDSC #27273), *UAS-Bpac* (BDSC #78788), *UAS-Cam2*.*1* (BDSC #6901), *UAS-CaMPARI* (BDSC #58761), *UAS-CaMPARI2* (BDSC #78316), *UAS-cAMPr* (Hackley et al., 2018), *UAS-Chrimson* (BDSC #55135), *UAS-Epac1-camps* (BDSC #25407), *UAS-GCaMP6f* (BDSC #42747), *UAS-GCaMP6s* (BDSC #42749), *UAS-GFP* (BDSC #32184), *UAS-GtACR1* (BDSC #92983), *UAS-myrGFP* (BDSC #32197), *UAS-nSyb-GFP* (BDSC #6921), *UAS-Piezo* (BDSC #78336), *UAS-RFP* (BDSC #27398), *UAS-Trhn-RNAi* (*Trhn-RNAi-1*, (Albin et al., 2015)), *UAS-TrpA1* (BDSC #26263), *VGlut-GAL80* (BDSC #58448).

#### Other lines

*OrgR* (BDSC #5), *Piezo*^*KO*^ (BDSC #58770).

### Construction of plasmids and generation of *lexAop2-CaMPARI* transgenic fly line

Standard molecular biology methods were used and constructs were sequence verified prior to microinjection into fly embryos. Restriction enzymes and T4 DNA ligase were from New England Biolabs. PCR amplifications were performed with Q5 polymerase (New England Biolabs).

First *CaMPARI* coding sequence was PCR amplified from plasmid *pcDNA3-CaMPARI* (gift from Loren Looger & Eric Schreiter, Addgene plasmid #60421) (Fosque et al., 2015) with primers 5’-GTCGACCATGCTGCAGAACGAGCTT-3’ and 5’ CTGATCAGCGAGCTCTAGCAT-3’. The PCR product was subcloned into *pCRII-TOPO* vector (Invitrogen) resulting in plasmid *TOPO-CaMPARI*. Then, *myrGFP* from *pJFRC19-13XLexAop2-IVS-myr::GFP* (gift from Gerald Rubin, Addgene #26224) (Pfeiffer et al., 2010) was removed by *Xho*I/*Xba*I digest and replaced with *Sal*I/*Xba*I fragment from *TOPO-CaMPARI* harboring *CaMPARI* coding sequence, generating plasmid *pIZ-CaMPARI*. Plasmid microinjections to generate two *lexAop-CaMPARI* fly lines (*P{y*^*+t7*.*7*^ *w*^*+mC*^*=13XLexAop2-CaMPARI}attP40* and *PBac{y*^*+*^ *w*^*+mC*^*=13XLexAop2-CaMPARI}VK00027*) were performed by BestGene Incorporated.

### Dissection of semi-intact larva

Feeding 3^rd^ instar larvae were dissected in petri dishes coated with a two-component silicone elastomer (Elastosil RT 601, Wacker Chemical Corporation). Larvae were pinned down dorsal side up at the posterior and anterior end using sharp-etched tungsten needles (∅ 77 μm). Larva was cut open longitudinally along the dorsal midline and thereafter the cuticle was cut transversely below the CPS with a micro scissors (15000-08, Fine Science Tools). Interior organs like fat body, trachea or salivary glands were removed except for the CNS and CPS including the associated pharyngeal nerves and digestive tract up to the anterior midgut. This standard preparation of the *Drosophila* larva was used in all experiments which involve dissection of larvae, in the following termed semi-intact preparation. Any further dissections are documented separately in the individual experimental sections.

### Immunohistochemistry

Dissected larval brains were fixed for 1 h in paraformaldehyde (4 %) in 1× phosphate-buffered saline (PBS), rinsed three times (20 min) with 1 % PBS-T (1 % Triton X-100 in 1× PBS), and blocked in 1 % PBS-T containing 5 % normal goat serum (ThermoFisher) for 2 h. Primary antibody was added to the solution (for concentrations, see below). Brains rotated 2 nights at 4 °C. On the third day, after removing the primary antibody, larval brains were washed three times (20 min) with 1 % PBS-T, and a secondary antibody was applied. Brains rotated two nights at 4 °C. After three times washing (20 min) with 1 % PBS-T, brains were dehydrated and cleared through an ethanol-xylene series and mounted in DPX Mountant (Sigma-Aldrich). Imaging was carried out using a Zeiss LSM 780 confocal microscope with LCI Plan-Neofluar 25× / 0.8 Imm Korr DIC M27 or Plan-Apochromat 63× / 1.4 Imm DIC objective (oil). For antibody staining of the *driver > GFP/myrGFP/nSyb-GFP*, the primary antibody was anti-GFP (1:500, chicken, Abcam, ab13970). Secondary antibody was anti-chicken Alexa Fluor 488 (1:500, goat, Invitrogen, A-11039). For 5-HT/Trhn staining, primary antibodies were anti-5-HT (1:1000, rabbit, Sigma, Lot #033M4805) and anti-Trhn (1:250, guinea pig, generated by Thermo, immunogen sequence: DSFEEAKEQMRAFAESIQR), secondary antibodies were anti-rabbit Alexa Fluor 633 (1:500, goat, Invitrogen, A-21071) and anti-guinea pig Alexa Fluor 633 (1:500, goat, Invitrogen, A-21105). For VGlut staining, primary antibody was anti-VGlut (1:1000, rabbit, gift from Hermann Aberle). The secondary antibody was anti-rabbit Alexa Fluor 633 (1:500, goat, Invitrogen, A-21052). For *5-HT7 > GFP* and *5-HT7 > GFP, VGlut-Gal80* staining, primary antibodies were anti-GFP (1:500, chicken, Abcam, ab13970) and anti-elav (1:500, mouse, DSHB, AB 528217). Secondary antibodies were anti-chicken Alexa Fluor 488 (1:500, goat, Invitrogen, A-11039) and anti-mouse Alexa Fluor 633 (1:500, goat, Invitrogen, A-21052). For background/neuropil staining, primary antibody was anti-22C10 (1:500, mouse, DSHB, AB528403). 22C10 was deposited to the DSHB by Seymour Benzer and Nansi Colley. Secondary antibody was anti-mouse Alexa Fluor 405 (1:500, goat, Invitrogen, A-31553). For serotonin receptor expression analysis following primary antibodies were used: anti-GFP (1:500, chicken, Abcam, ab13970), anti-sNPF (1:1000, rabbit, gift from Jan Veenstra) or anti-pros (1:500, mouse, DSHB, AB 528440). Accordingly, secondary antibodies were anti-chicken Alexa Fluor 488 (1:500, goat, Invitrogen, A-11039) and anti-rabbit Alexa Fluor 633 (1:500, goat, Invitrogen, A-21071) or anti-mouse Alexa Fluor 633 (1:500, goat, Invitrogen, A-21052). For occasional F-actin staining, we used the conjugated fluorescent Phalloidin-TRITC (1:1000, ThermoFisher).

### Functional Imaging

For Calcium-imaging by an integrator, we used CaMPARI *(Fosque et al., 2015)* and CaMPARI2 (Moeyaert et al., 2018). In the experiments with *peb > CaMPARI2* and *Piezo*^*Gal4*.*KI*^ *> CaMPARI2* a starved larva (≥ 30 min starvation time) was placed on either a water agar plate (non-fed condition) or a water agar plate coated with yeast (fed condition). 405 nm UV light (M405L2, Thorlabs) connected to a LED controller (LEDD1B, Thorlabs) was positioned 12 cm above the larva and illuminated at max intensity for 2 min. Afterwards the larval esophagus was dissected and put onto a poly-L-lysine-coated coverslip and covered with 1× PBS for imaging at low Ca^2+^ conditions. EG neurons with their dendrites which cover the esophagus were imaged. For the CaMPARI experiments of Se0 neurons, we used *Trhn/Se0*_*ens*_*/Se0*_*ph*_ *> CaMPARI*. In gustatory experiments the larva was placed in a well of a Terasaki plate (Cat.-No. 659180, Greiner) filled with 20 μl 10 % yeast-/ 1 M fructose-/ 20 mM caffeine-/ 10 mM denatonium-/ 2 M NaCl-solution (solved in tap water)/ tap water. Or wells were filled with 20 μl 0.5 M fructose solution at different hydroxypropyl cellulose (HPC) concentration for mechanical stimulation. 405 nm UV light was positioned 12 cm above the larva and illuminated at max intensity for 30 s after a 2 min rest period. Afterwards the larval brain was dissected and placed onto a poly-L-lysine-coated coverslip and covered with 1× PBS for imaging at low Ca^2+^ conditions. The SEZ region with the Se0 neurons was imaged. For the CaMPARI experiments of Se0 neurons while activating EG_post_ neurons, we used the genotype *Piezo*^*III*^*-Gal4 > UAS-TrpA1; Trhn-LexA > lexAop-CaMPARI*. Prior to the experiment larvae were starved (≥ 30 min starvation time) and transferred on either a water agar plate or water agar plate coated with yeast for 2 min. Afterwards the larvae were dissected. A custom-made heating device for local thermal application was positioned close to the EG_post_ neurons without covering the CNS. A 405 nm UV light was positioned 6 cm above the semi-intact larvae. Simultaneously the larvae were illuminated at max intensity and thermal stimulus of 18 °C (no activation) or 32 °C (activation) was applied to EG_post_ neurons for 2 min. After this time period the brain was dissected and positioned on a poly-L-lysine-coated coverslip with 1× PBS for imaging at low Ca^2+^ conditions. The Se0 neurons in the SEZ were imaged. All images were acquired using a ZEISS LSM 780 Laser scanning microscope with LCI Plan-Neofluar 25 × / 0.8 Imm Korr DIC M27. For quantification, green to red ratios of single cells were analyzed with a custom-made script for FIJI (ImageJ), and the mean per animal was calculated (each cell was analyzed and a mean calculated). Obtained data was then statistically analyzed and plotted with SigmaPlot (version 12) software using the Mann–Whitney rank-sum test.

For GCaMP-recordings, the genotypes *Mef2 > GCaMP6f, Mef2 > GCaMP6s,RFP* and *30F10 > GCaMP6f* were used. To record the ERM or neural activity of ERM-MNs in the HCG, dissected foregut and ENS of a larva based on semi-intact preparation was positioned on a poly-L-lysine-coated coverslip and covered with 18 μl saline (Rohrbough and Broadie, 2002). Images were acquired with a Zeiss LSM 780 laser scanning microscope as time series with a scan speed of approximately 50 ms (∼20 Hz) using a Zeiss LCI Plan-Neofluar 25× / 0.8 Imm Korr DIC M27 objective dipped in the saline solution. After 3 min initial recording 2 μl 10^−6^ M 5-HT solution was added to obtain a final experimental concentration of 10^−7^ M 5-HT. For quantification, recordings were analyzed using the software Zen 2012 (Zeiss) and a custom-made script for Spike2 (Cambridge Electronic Design) to measure cycle frequency and completion rate for each experiment. The completion rate is the percentage of induced contraction waves in the initiation zone which successfully conveyed into the propagation zone representing a complete esophageal peristalsis.

For cAMP-level measurement using the genotypes *OK371 > Epac1-camps*, larva was dissected and a semi-intact preparation was positioned on a poly-L-lysine-coated coverslip with 20 μl saline (Rohrbough and Broadie, 2002) or fresh prepared 10^−7^ M 5-HT solution (solved in saline). 3 min after 5-HT application the ERM-MNs in the HCG were scanned. All images were acquired using a ZEISS LSM 780 Laser scanning microscope with LCI Plan-Neofluar 25 × / 0.8 Imm Korr DIC M27. For quantification, cyan fluorescence to yellow fluorescence red ratios for individual cells were measured with a custom-made script for FIJI (ImageJ), and the mean was calculated per animal (each cell was analyzed and a mean build). Animal means (including standard error of mean) were then analyzed and plotted with SigmaPlot (version 12) software.

For cAMP-level measurement using the genotypes *VGlut > cAMPr*, larva was dissected and a semi-intact preparation was positioned on a poly-L-lysine-coated coverslip covered with 20 μl saline (Rohrbough and Broadie, 2002) or fresh prepared 10^−7^ M 5-HT solution (solved in saline). Imaging time series lasted 600 s in which each 30s an image was scanned. All images were acquired using a ZEISS LSM 780 Laser scanning microscope with LCI Plan-Neofluar 25× / 0.8 Imm Korr DIC M27. For quantification, green fluorescence intensity for individual cells were measured with a custom-made script for FIJI (ImageJ). Mean of all analyzed cells (including standard error of mean) for each time point was analyzed and plotted with SigmaPlot (version 12) software.

### Electrophysiological recordings

For extracellular recordings, semi-intact larva preparation was used to expose the pharyngeal nerve, e.g. antennal nerve (AN), the eye-antennal disk was removed. The nerve was insulated with a paraffin-petroleum jelly mixture. Neural activity was measured using custom made silver wire electrodes connected to an amplifier/signal conditioner system (Model MA 102&103, Neuroscience Electronics Laboratory, University of Cologne). All recorded signals were amplified (5,000 fold) and filtered (0.1– 3 kHz). Recordings were sampled at 20 kHz. Data was acquired with Power 1401 mk II A/D board and Spike2 software (Cambridge Electronic Design). To measure only afferent signals, the CNS was removed after establishing the nerve recording.

### Optogenetic manipulation

For optogenetic manipulation of neuronal activity, two effector lines *UAS-GtACR1* (Mohammad et al., 2017) for inhibition and *UAS-Chrimson* (Oda et al., 2018) for excitation were used in the respective experiments. As mentioned, larvae were bred on standard cornmeal medium containing 150μM all-trans retinal. Activation of these light-gated channels was induced by illumination at 530 nm (Chrimson) with a laser LED (M530L3, Thorlabs) or 625 nm (GtACR1) with a laser LED (M625L3, Thorlabs). In electrophysiological experiments the laser LEDs were attached to an optical fiber system (Thorlabs) to stimulate/inhibit specific regions of the ENS. For behavioral assays, the laser LED was mounted onto a collimated lens to illuminate the experimental area. LEDs were controlled via an A/D board (Power 1401 mk II, Cambridge Electronic Design) which was connected to voltage-controlled LED power supply (LEDD1B, Thorlabs). Stimulus timing and duration were set by protocols in Spike2 software (Cambridge Electronic Design). All optogenetic experiments were performed in darkness.

### Thermogenetic manipulation

For thermogenetic manipulation of neuronal activity, *UAS-TrpA1* (Pulver et al., 2009) for excitation was used in respective experiments. Local thermal stimulus was applied with a custom-made heating device controlled by an A/D board (Schoofs et al., 2014). The local temperature of a tissue was shifted to 18 °C for non-activating or shifted to 32 °C for activating the respective TrpA1 expressing neuronal tissue.

### Behavioral assays

For short-term food intake assay, only feeding third instar larvae (96±2 h) were used. Apple juice agar plates were prepared with a spot of colored yeast paste in the middle of the plate. Before the experiment plates were placed RT for 2 h. After 30 min starvation, five larvae were transferred on top of colored yeast paste for 20 min. Subsequently, larvae were transferred into a cell strainer and washed with 65 °C hot water. Larvae were then transferred onto glass slides for photo documentation and analyzed with FIJI software (ImageJ) by a custom written analysis macro, which determined the percentage of the colored surface of the intestinal system compared to body surface of the larva.

To monitor the motility of the larval foregut, videos of semi-intact larvae were recorded using a camera (Quickcam Pro 9000, Logitech or C1 Pro, Kurokesu) mounted to a microscope (Stemi-2000C, Zeiss). Dark-field microscopy was used to improve visibility. VirtulDub or Spike2 was used as capturing software. For pharmacological experiments, a well was filled with 45 μl of saline solution and additional 5 μl of varying 5-HT solutions. In control experiments 5 μl of saline solution were instead applied. Optogenetic experiments were performed in a silicone elastomer-coated petri dish filled with saline solution. Recorded videos were manually analyzed three times to determine the mean activity of ERM as peristaltic waves per minute. Only complete peristaltic waves from pharynx to proventriculus counted as activity.

### EM reconstruction

Neuron reconstruction was done on a STEM (scanning transmission electron microscopy) volume of a whole first instar larva; the technical details of its generation will be described separately by Cardona and colleagues (Peale et al., in prep.). All reconstructions were made in a modified version of CATMAID (http://www.catmaid.org, (Saalfeld et al., 2009)). For reconstructing a neuron, a specific neurite in a section of the electron microscopy data set was identified and a neuronal 3D skeleton including the synaptic active zones and synaptic partners was manually generated. We identified all enteric neurons, Se0 neurons and specific pharyngeal neurons by reconstruction of all axons passing through the frontal nerve junction (FNJ) originating either in the CNS or ENS. We reconstructed all neurons to completion (tracing 100% and at least 95% reviewed).

For the Se0 neurons, as modulatory output neurons (Miroschnikow et al., 2018), all membrane fusion sides of CCVs (clear core vesicles) were marked as connectors without synaptic partners. To determine the putative downstream targets of Se0 neurons all tissues innervated by the ENS, e.g. foregut, midgut, ring gland and garland cells were reconstructed. For identified motor neurons, the innervated muscle was reconstructed and the neuromuscular junctions (NMJs) marked as connectors targeting the respective muscle. For identified neuropeptidergic neurons, the number of cytoplasmic DCVs (dense core vesicles) was determined.

### Statistical Information

All statistical analyses were carried out in SigmaPlot (12.0) or PRISM 9.0 (GraphPad). All performed statistical tests, number of replicates and statistical significance values of represented data are reported in the corresponding figure, figure legends and in the corresponding section of the Materials and Methods. In all boxplots shown in Figure and Extended Data Figure, the solid line depicts the median; the upper and lower boundary of the box depict the first and third quantiles of the data set, respectively. Whiskers indicate 5% and 95% confidence level. Individual data points of the box plots are included as circles. All measurements were taken from distinct samples.

## Supplemental figures

**Figure S1:**
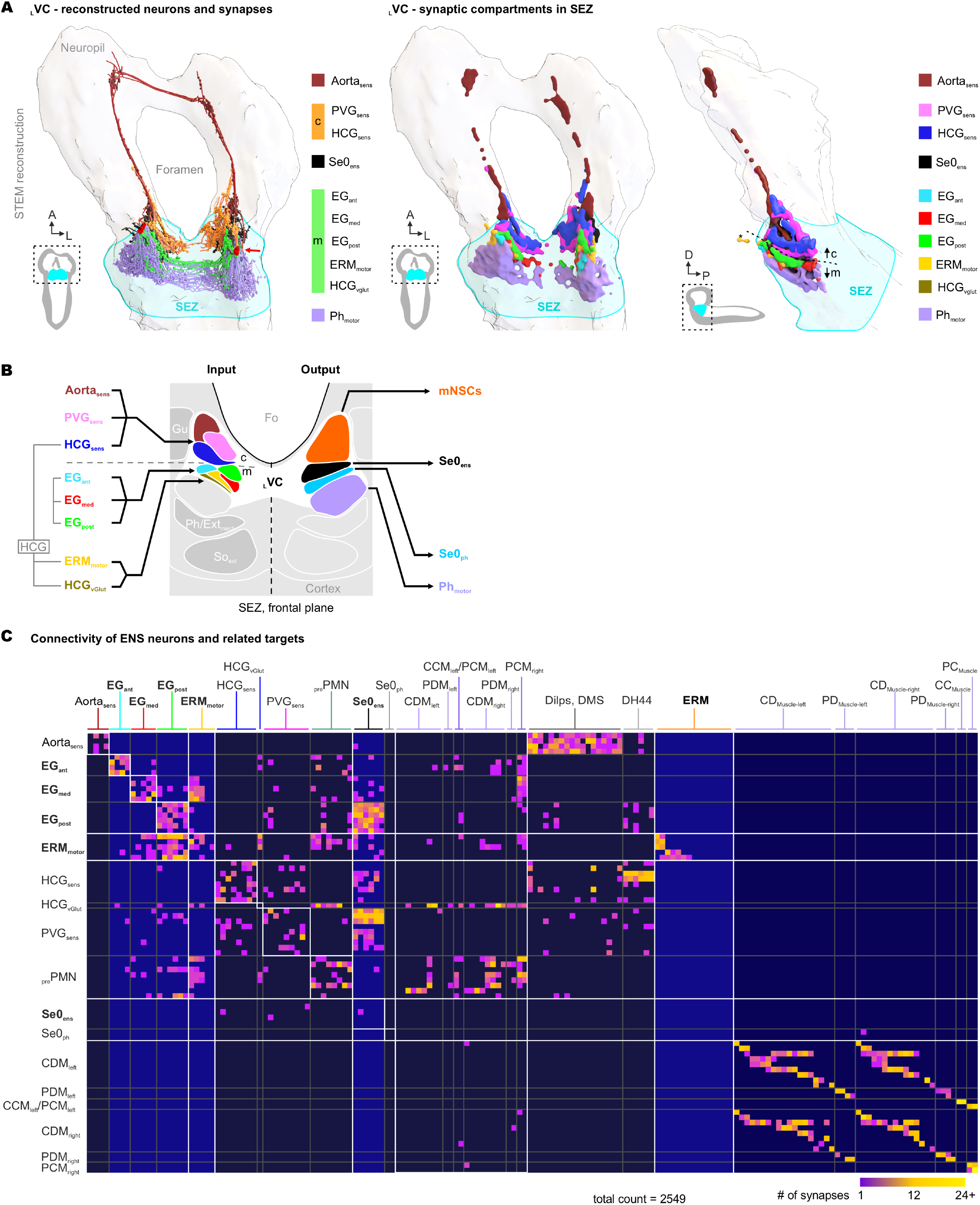
Reconstructed CNS-ENS connectome of *Drosophila* larva. **(A)** Left: Three-dimensional illustration of the reconstructed neurons including the synapses for the larval vagus complex (_L_VC). Right: Three-dimensional illustration of the relevant synaptic compartments. Note, for the aorta, we were able to identify two sensory neurons per side (Aorta_sens_) with their receptive field on the aorta which have synaptic connections to the protocerebrum and the neurosecretory system (Hückesfeld et al., 2021; Schlegel et al., 2016). **(B)** Heat-map showing the connectivity of ENS neurons and related targets, like the pharyngeal motor system. **(C)** Illustration of the synaptic input (sensory) and synaptic output (motor/modulatory) compartments of the larval vagus center (_L_VC) in the subesophageal zone (SEZ). **Abbr**.: Aorta_sens_ - sensory neurons of the aorta, CCM_left/right_ – cibarial constrictor muscle (left/right), CDM_left/right_ – cibarial dilator muscle (left/right), DH44 – diuretic hormone 44, Dilps – *Drosophila* insulin-like peptide, DMS - drosomyosuppresin, EG_ant/med/post_ – esophageal ganglion (anterior, medial, posterior), ERM – esophageal ring musculature, ERM_motor_ – ERM motor neuron, HCG_vGlut_ – VGlut-positive HCG neuron, HCG_sens_ – sensory HCG neuron, mNSC - medial neurosecretory cells, PCM_left/right_ – pharyngeal constrictor muscle (left/right), PDM_left/right_ – pharyngeal dilator muscle (left/right), Ph_motor_ – pharyngeal motor neuron, _pre_PMN – pharyngeal premotor neuron, PVG_sens_ – sensory PVG neuron, Se0_ens_ – enteric Se0 neurons, Se0_ph_ – pharyngeal Se0 neuron.

**Figure S2.**
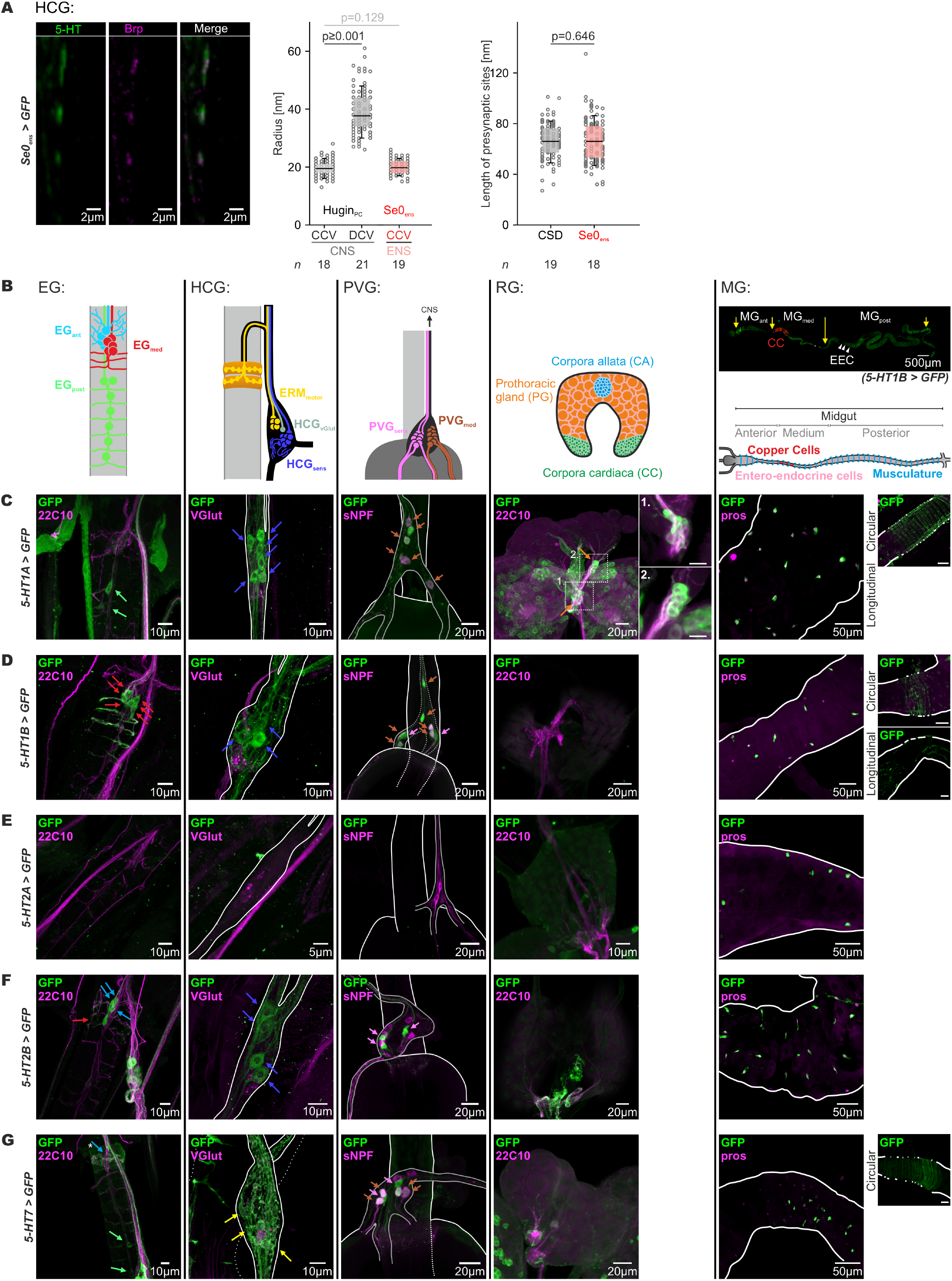
Peripheral presynaptic sites of Se0_ens_ neurons and serotonin receptor expression analysis in the ENS. **(A)** Left: antibody staining of *29H01 > GFP* against serotonin (5-HT) and *Bruchpilot* (*Brp*) indicating peripheral presynaptic sites of serotonergic Se0_ens_ neurons in the HCG. Middle: box plots showing the radius of CCV (clear core vesicles) and DCV (dense core vesicles) for Hugin_PC_ neurons (grey) in the CNS compared to the radius of CCV of Se0_ens_ neurons (red) in the ENS. Note that there is no significant difference between the CCVs of Hugin_PC_ and Se0_ens_. Right: comparison between the length of peripheral presynaptic sites of serotonergic Se0_ens_ neurons and central presynaptic sites of the serotonergic CSD neuron. There is no significant difference in length between the central and peripheral presynaptic sites. Data obtained from brain STEM volume. **(B)** Schematic drawings of the ganglions of the ENS including their different neuron types (colored), larval endocrine organ (rind gland) and midgut which were analyzed for their expression of serotonin receptors. **(C-G)** Fluorescence images show the GFP-expression for the serotonin receptors 5-HT1A (**C**), 5-HT1B (**D**), 5-HT2A (**E**), 5-HT2B (**F**) and 5-HT7 (**G**) in ENS, RG and MG of *Drosophila melanogaster*. To identify the different cell types in the ENS or MG additional antibody stainings against VGlut (motor neurons), sNPF (modulatory neurons), pros (entero-endocrine cells) or 22c10 (neurites) were included. Colored arrows mark identified cell bodies of neurons. **Abbr**.: EG_ant/med/post_ – esophageal ganglion (anterior, medial, posterior), ERM_motor_ – ERM motor neuron, HCG – hypocerebral ganglion, HCG_sens_ HCG sensory neuron, HCG_vGlut_ – VGlut-positive HCG neuron, MG - midgut, PVG – proventricular ganglion, PVG_mod_ –PVG modulatory (local) neuron, PVG_sens_ –PVG sensory neuron, RG – ring gland.

**Figure S3.**
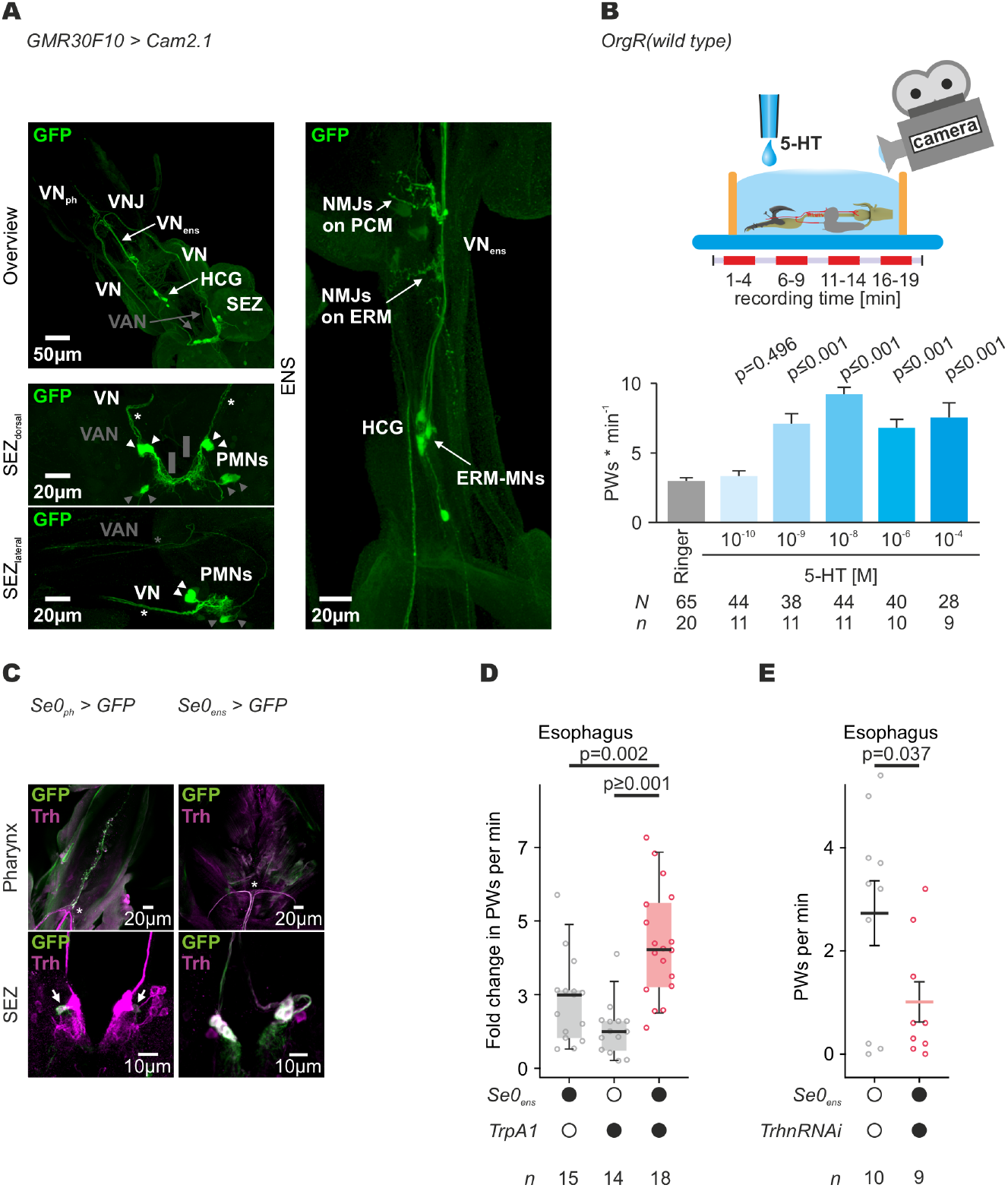
Expression pattern of GMR30F10-Ga4 and effect of serotonin on esophageal ring muscle (ERM) motor system. **(A)** GMR30F10 > Cam2.1 shows expression three different types of feeding-related motor neuron types: 1) ERM_motor_ which are located in HCG, 2)PMNs (marked by white arrowheads) projecting through the (VN (marked by white asterisks) which innervate via VN_ph_ the CDM and via VN_ens_ the PCM and 3) PMNs (marked by gray arrowheads) projecting through the VAN (marked by gray asteriks) which innervate the PDM. **(B)** Top: Schematic drawing of the experimental setup to monitor esophageal motility in semi-intact larva. Semi-intact larva was placed in an experimental well filled with 50 μl 5-HT/saline solution. Larva was not fixated to avoid mechanoreceptive activation. Esophageal movements were analyzed in successive video-recordings. Bottom: bar plot shows the dose-dependent increase peristaltic waves (PWs) per minute after bath application of 5-HT (data shows mean and ±SE). **(C)** Antibody staining of Se0_ph_ > GFP and Se0_ens_ > GFP. The Gal4 driver lines show distinct expressions in the two Se0 subclusters. The one Se0_ph_ neuron projects to pharynx via VN_ph_ (indicated by asterisk). The three Se0_ens_ neurons project into the entire ENS via VN_ens_ (indicated by asterisk). **(D)** Box plot shows that activating Se0_ens_ neurons accelerates peristaltic waves (PWs) per minute compared to controls. Performed significance test: Mann-Whitney rank sum test. **(E)** Graph indicates that blocking serotonin synthesis in Se0_ens_ neurons by RNAi against tryptophan hydroxylase (*Trhn*) reduces PWs per min. Performed significance test: Mann-Whitney rank sum test. **Abbr**.: ERM - esophageal ring musculature, ERM_motor_ - ERM motor neurons, HCG - hypocerebral ganglion, NMJ -neuromuscular junction, PCM - pharyngeal constrictor musculature, PMN - pharyngeal motor neuron, SEZ - subesophageal zone, VAN - ventral arm nerve, VN - vagus nerve, VN_ens_ - enteric vagus nerve, VN_ph_ - pharyngeal vagus nerve, VNJ - vagus nerve junction.

**Figure S4.**
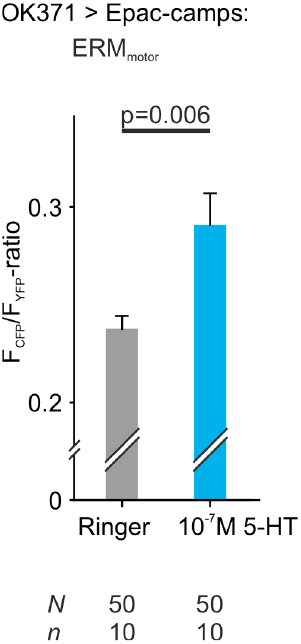
Application of serotonin increase intracellular cAMP-level in ERM_motor_. The cAMP-reporter, Epac1-camps, showed an increased cAMP-level in ERM_motor_ after serotonin treatment using the Gal4 driver line *OK371*. Data shows mean ±STD. Performed significance test: Mann-Whitney rank sum test. **Abbr**.: cAMP – cyclic adenosine monophosphate, ERM_motor_ - esophageal ring musculature motor neuron.

**Figure S5.**
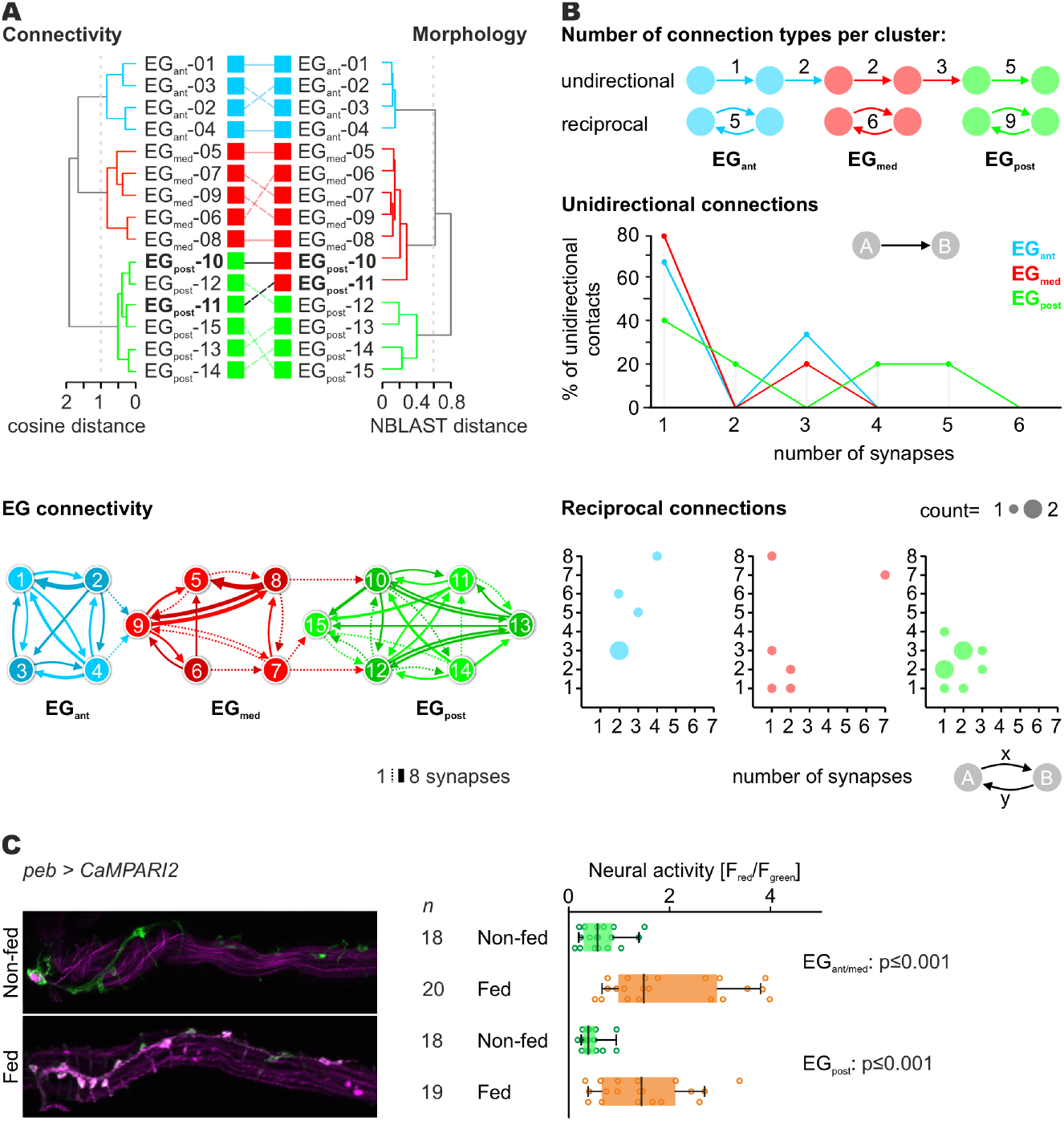
EG neuron connectivity and sensory perception upon food intake. **(A)** Top: Cluster analysis of EG neurons by neuronal morphology and synaptic connectivity indicates that the EG consists of three entities. Bottom: EG connectivity diagram showing the synaptic connections of each individual EG neuron. Line thickness represents the number of synaptic connections. Arrows with dashed lines mark synaptic connections with only one synapse. **(B)** Top: Comparison between unidirectional and reciprocal connections of the three EG clusters. Middle: graph shows the percentage of unidirectional connections in relation to the number of synapses for each EG cluster. Bottom: dot plots show the number of synapses for each neuronal partner in a reciprocal connection for the EG clusters. Diameter of the dots represents the quantity of a reciprocal connection. **(C)** Representative images of esophageal ganglion show the increased neural activity upon food passage (top). Box plot shows the significantly increased neural activity between non-fed and fed state of EG_ant/med_ and EG_post_. Performed significance test: Mann-Whitney rank sum test. **Abbr**.: EG_ant_ - anterior esophageal ganglion, EG_med_ - medial esophageal ganglion, EG_post_ - posterior esophageal ganglion,.

**Figure S6.**
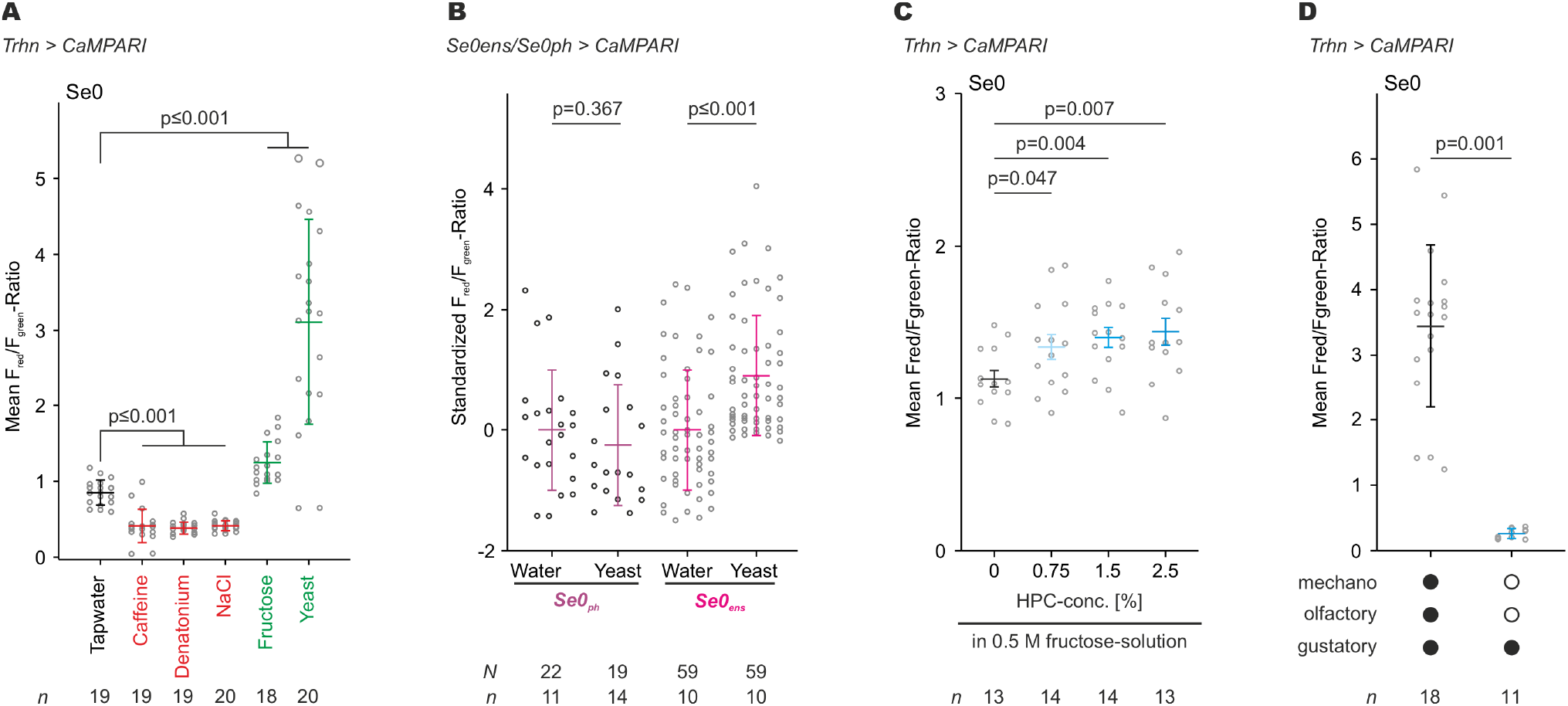
Se0_ens_ neurons respond to attractive nutrients. **(A)** CaMPARI-experiments showed that Se0 neurons respond with a decrease in neural activity after presentation/ingestion of aversive nutrients, e.g. caffeine, denatonium and NaCl (marked red). Otherwise ingestion of attractive nutrients, like fructose and yeast, resulted in increased neural activity (marked green). Performed significance test: Mann-Whitney rank sum test. **(B)** CaMPARI-experiments of Se0_ph_ and Se0_ens_ neurons revealed that only Se0_ens_ neurons showed increased neural activity upon ingestion of attractive nutrients but not Se0_ph_ neurons. Data is shown as scatter plots (gray) including mean (colored line) and standard error (colored whiskers). Performed significance test: Mann-Whitney rank sum test. **(C)** Uptake of a 0.5 M fructose solution with different viscosity by addition of hydroxypropyl cellulose (HPC) resulted in increased neural activity of Se0 neurons. Performed significance test: student’s t-test (c). **(D)** Limiting the yeast experience in larva to gustatory perception by gluing the mouth opening resulted in a decreased neural activity of the Se0 neurons compared to free-behaving larva. Performed significance test: Mann-Whitney rank sumtest. **Abbr**.: Se0 - subesophageal cluster 0 of serotonergic neurons, Se0_ens_ - enteric Se0 neurons, Se0_ph_ - pharyngeal Se0 neurons.

**Figure S7.**
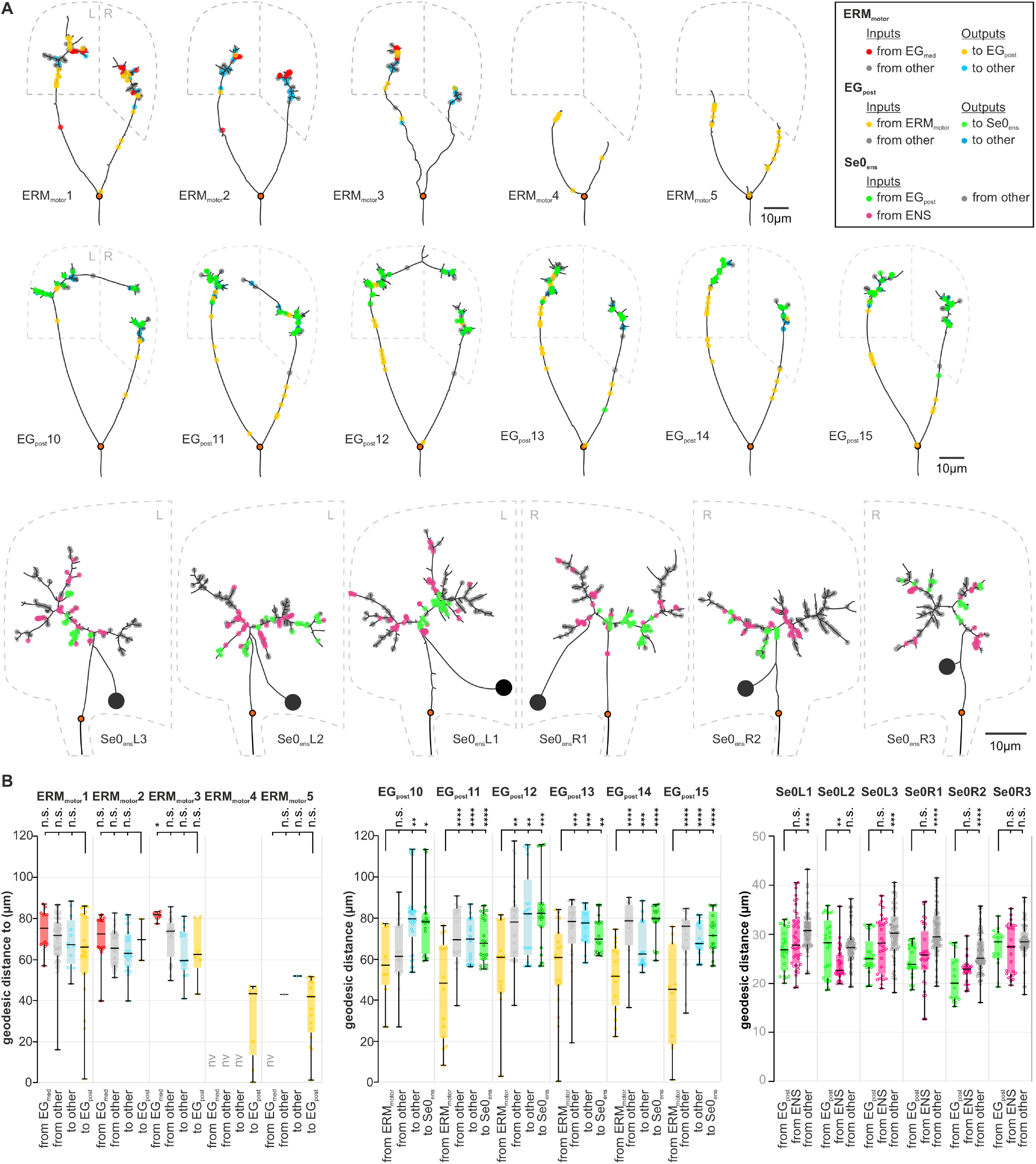
Synapse topology of the swallowing circuit at single cell level. **(A)** Two-dimensional dendrograms of neurons involved sensorimotor circuit for swallowing: **(B)** Spatial synapse analysis for ERM_motor_1-5_r_, EG_post_ 10-15 and Se0_ens_ L/R1-3 at single cell level. Box plots show the geodesic distance of specific pre- and postsynaptic sites for the three neuronal components of the swallowing circuit. Origin for the geodesic distance measurement (orange circle in Figure S7A) is the vagus nerve junction (VNJ) for ERM_motor_/EG_post_ and nerve entry site for Se0_ens_. Performed statistical test: one-way ANOVA. **Abbr**.: ENS - enteric nervous system, ERM_motor_ - Esophageal ring musculature motor neuron, EG_post_ - posterior esophageal ganglion neuron, Se0_ens_ - enteric subesophageal cluster 0 neuron.

**Table S1.**
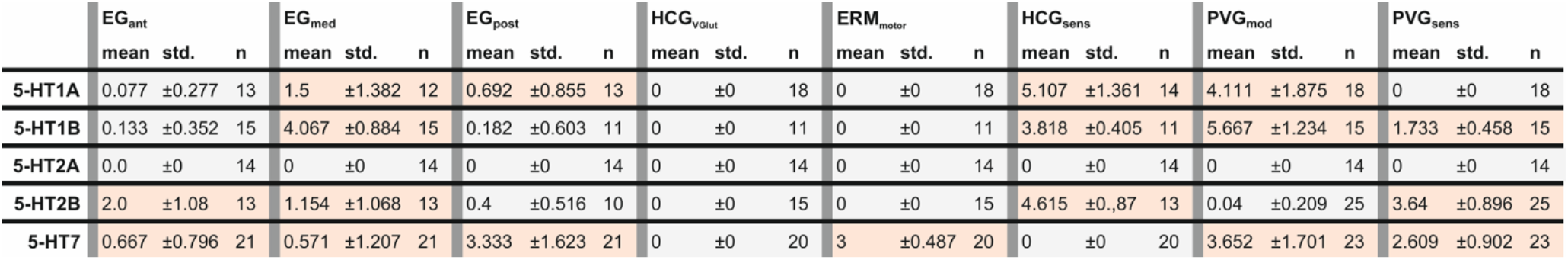
Expression-analysis of the enteric nervous system. Table shows the number of neurons expressing the serotonin (5-HT) receptor 1A, 1B, 2A, 2B and 7 based on immunohistochemical staining. Listed values are the mean, standard deviation (std.) and number of analyzed enteric nervous systems. Note, that for this analysis data from three different reporter lines were used (GFP, myrGFP and Cam2.1).

|  | EG<sub>ant</sub> |  |  | EG<sub>med</sub> |  |  | EG<sub>post</sub> |  |  | HCG<sub>VGlut</sub> |  |  | ERM<sub>motor</sub> |  |  | HCG<sub>sens</sub> |  |  | PVG<sub>mod</sub> |  |  | PVG<sub>sens</sub> |  |  |
| --- | --- | --- | --- | --- | --- | --- | --- | --- | --- | --- | --- | --- | --- | --- | --- | --- | --- | --- | --- | --- | --- | --- | --- | --- |
|  | mean | std. | n | mean | std. | n | mean | std. | n | mean | std. | n | mean | std. | n | mean | std. | n | mean | std. | n | mean | std. | n |
| 5-HT1A | 0.077 | ±0.277 | 13 | 1.5 | ±1.382 | 12 | 0.692 | ±0.855 | 13 | 0 | ±0 | 18 | 0 | ±0 | 18 | 5.107 | ±1.361 | 14 | 4.111 | ±1.875 | 18 | 0 | ±0 | 18 |
| 5-HT1B | 0.133 | ±0.352 | 15 | 4.067 | ±0.884 | 15 | 0.182 | ±0.603 | 11 | 0 | ±0 | 11 | 0 | ±0 | 11 | 3.818 | ±0.405 | 11 | 5.667 | ±1.234 | 15 | 1.733 | ±0.458 | 15 |
| 5-HT2A | 0.0 | ±0 | 14 | 0 | ±0 | 14 | 0 | ±0 | 14 | 0 | ±0 | 14 | 0 | ±0 | 14 | 0 | ±0 | 14 | 0 | ±0 | 14 | 0 | ±0 | 14 |
| 5-HT2B | 2.0 | ±1.08 | 13 | 1.154 | ±1.068 | 13 | 0.4 | ±0.516 | 10 | 0 | ±0 | 15 | 0 | ±0 | 15 | 4.615 | ±0.87 | 13 | 0.04 | ±0.209 | 25 | 3.64 | ±0.896 | 25 |
| 5-HT7 | 0.667 | ±0.796 | 21 | 0.571 | ±1.207 | 21 | 3.333 | ±1.623 | 21 | 0 | ±0 | 20 | 3 | ±0.487 | 20 | 0 | ±0 | 20 | 3.652 | ±1.701 | 23 | 2.609 | ±0.902 | 23 |

**Table S2.**
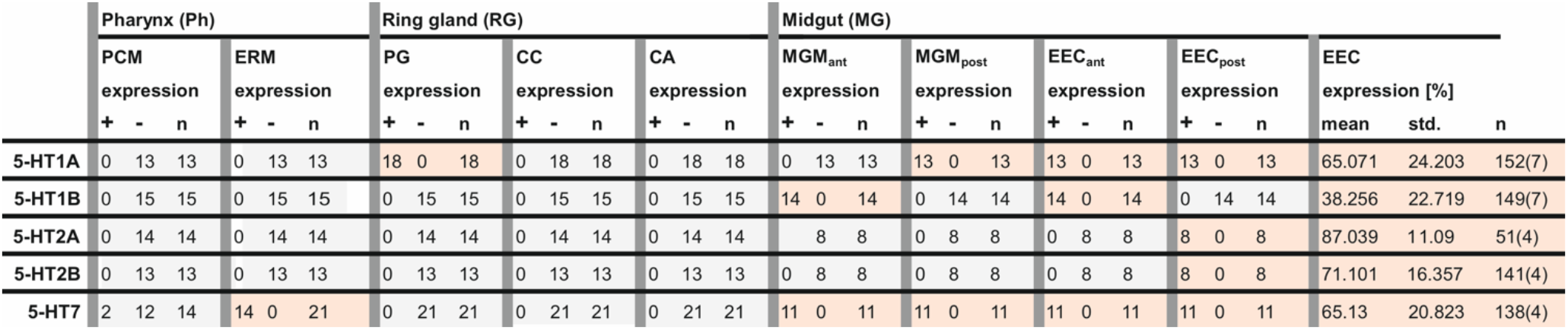
Expression-analysis of tissues associated with the enteric nervous system. Table shows the number of tissues expressing the serotonin (5-HT) receptor 1A, 1B, 2A, 2B and 7 based on immunohistochemical staining. Listed values display the number of 5-HT receptor expressing tissue (+), not 5-HT receptor expressing tissue (-) and total number of analyzed structures (n). Note, that for this analysis data from three different reporter lines were used (GFP, myrGFP and Cam2.1).

**Table S3.**
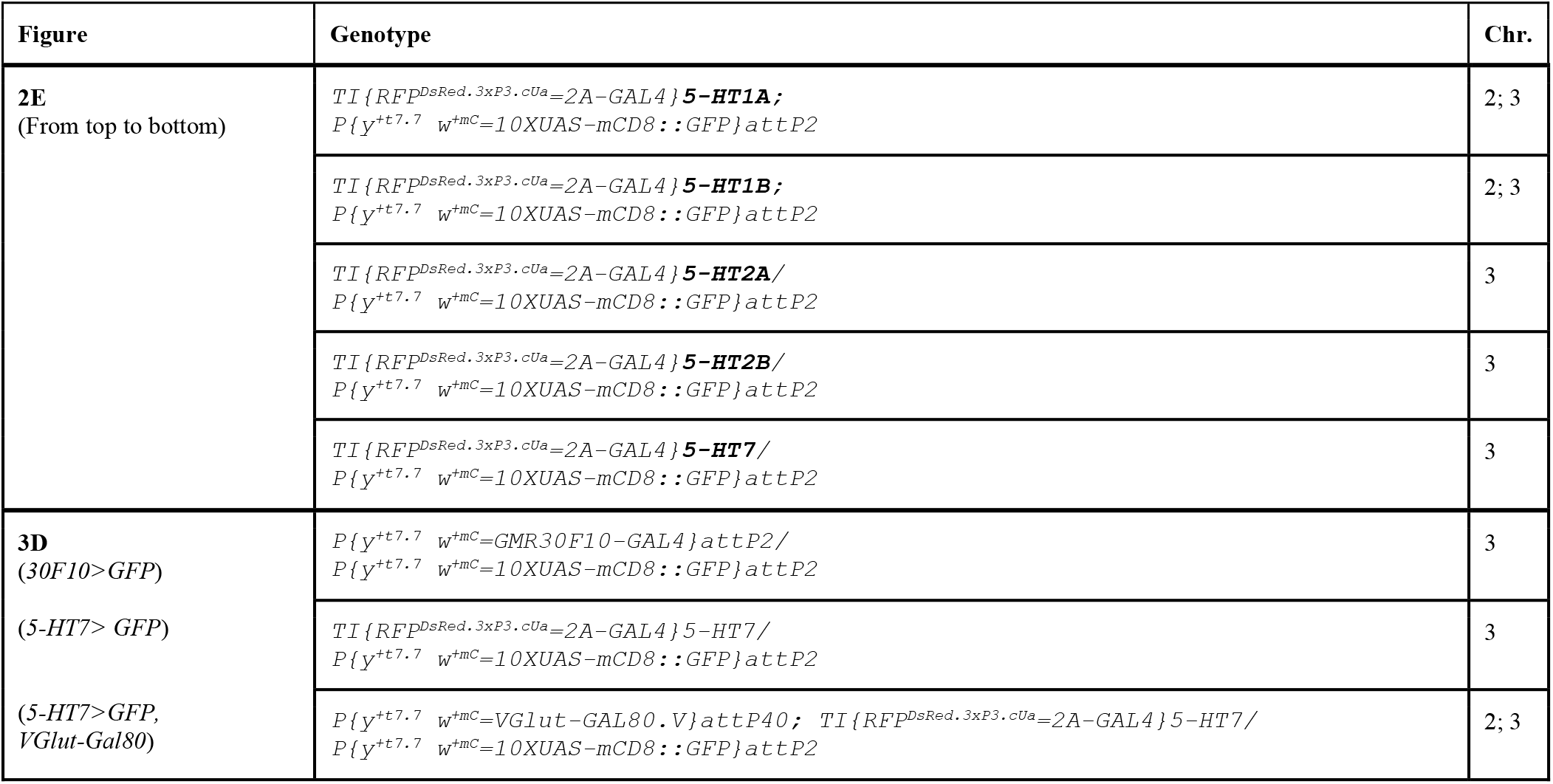
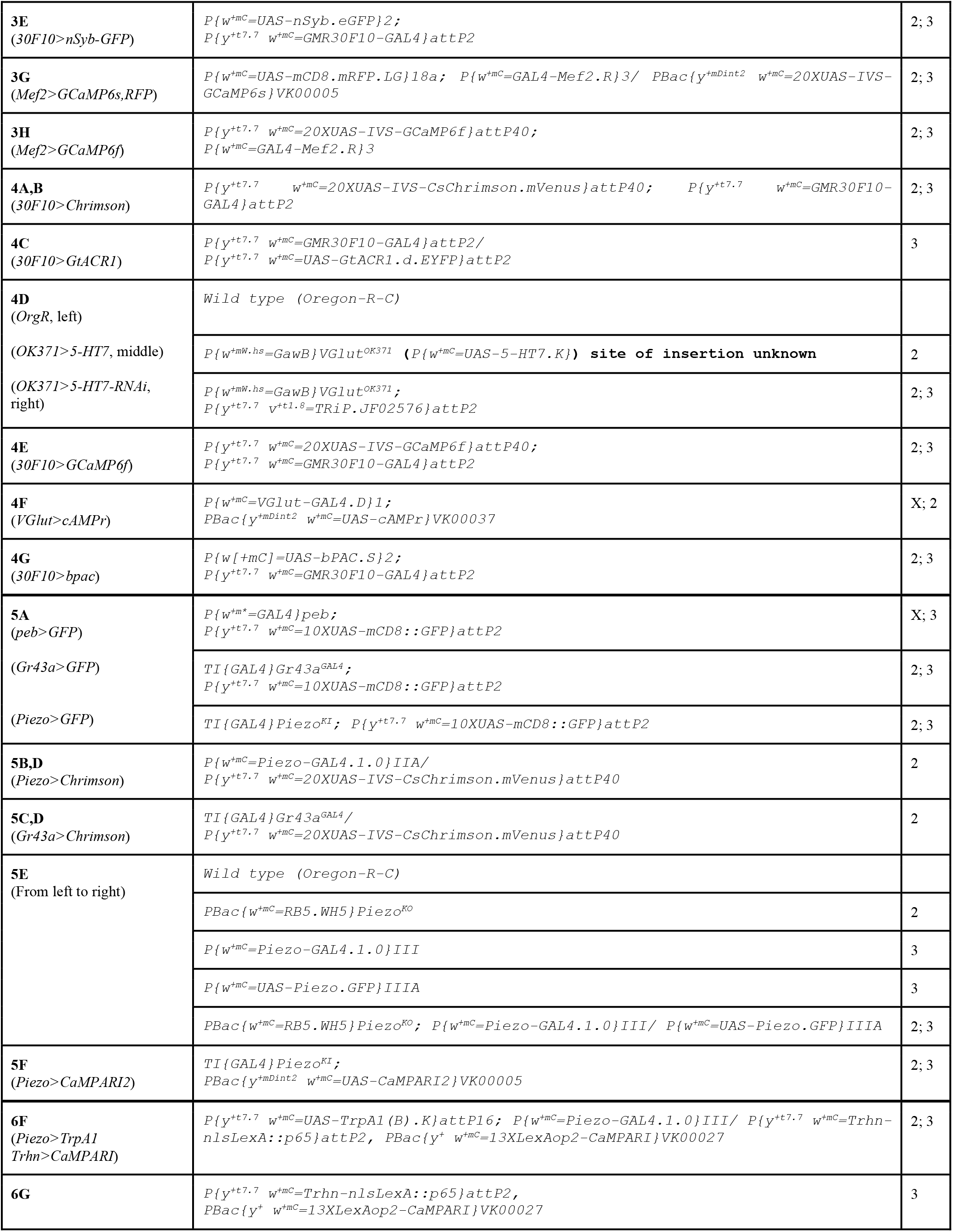
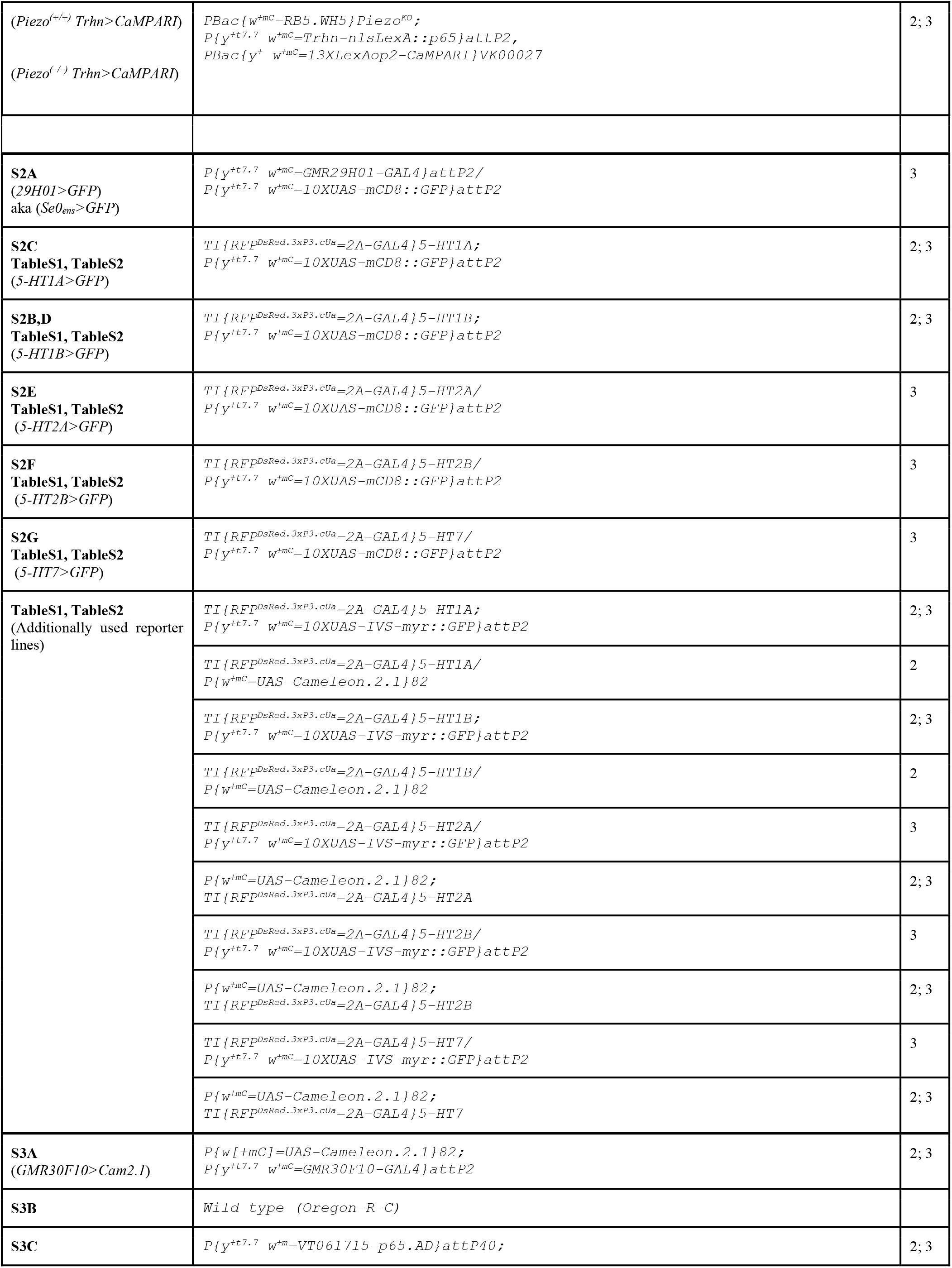
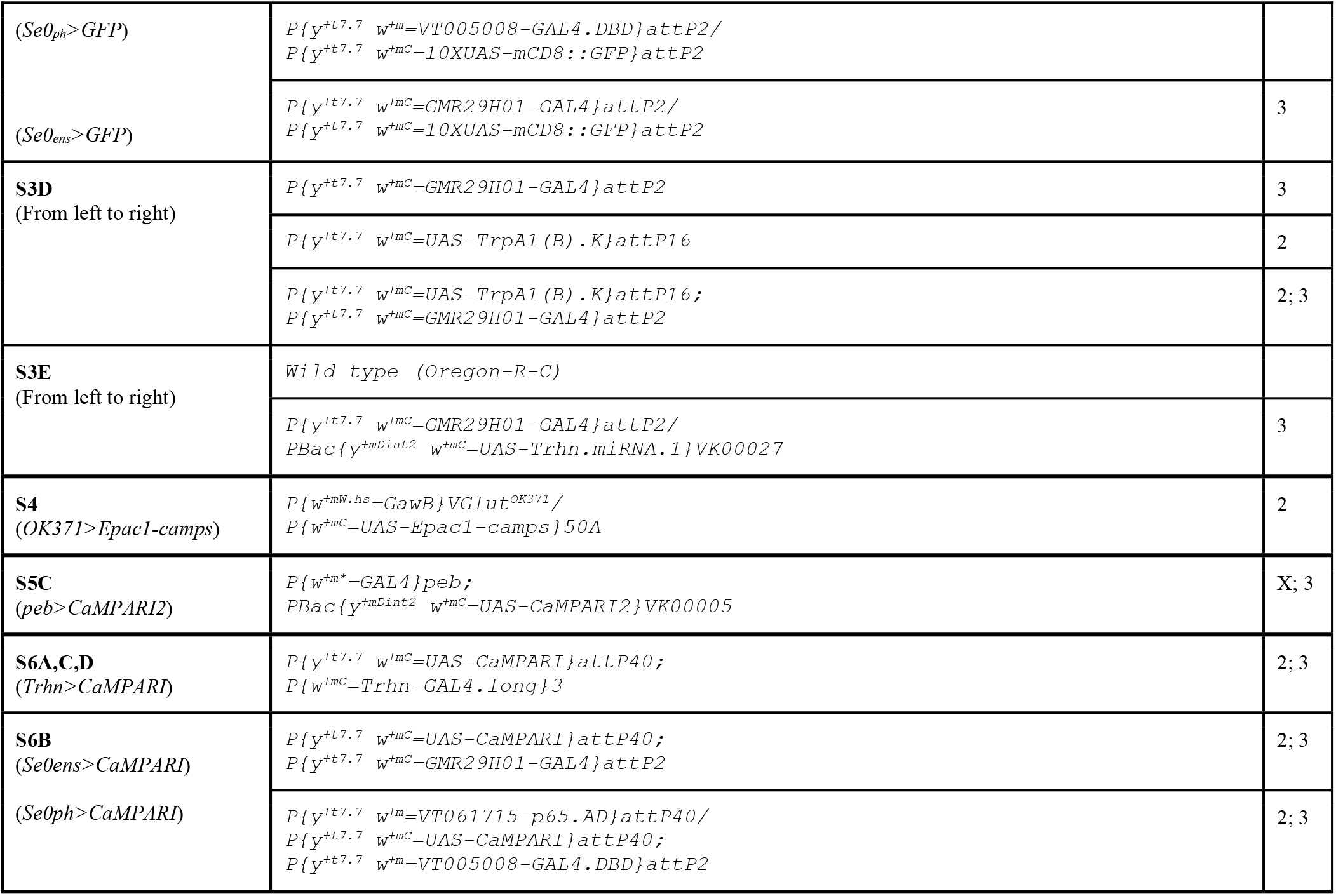
Genotypes of experimental flies.

